# Potential and action mechanism of favipiravir as an antiviral against Junin virus

**DOI:** 10.1101/2021.12.14.472716

**Authors:** Vahid Rajabali Zadeh, Shuzo Urata, Tosin Oladipo Afowowe, Jiro Yasuda

## Abstract

Favipiravir is a nucleoside analogue that inhibits the replication and transcription of a broad spectrum of RNA viruses, including pathogenic arenaviruses. In this study, we isolated a favipiravir-resistant mutant of Junin virus (JUNV), which is the causative agent of Argentine hemorrhagic fever, and analyzed the antiviral mechanism of favipiravir against JUNV. Two amino acid substitutions, N462D in the RNA-dependent RNA polymerase (RdRp) and A168T in the glycoprotein precursor GPC, were identified in the mutant. GPC-A168T substitution enhanced the efficiency of JUNV internalization, which explains the robust replication kinetics of the mutant in the virus growth analysis. Although RdRp-N462D substitution did not affect polymerase activity levels in a minigenome system, comparisons of RdRp error frequencies showed that the virus with RdRp-D462 possessed a significantly higher fidelity. We also provided experimental evidence for the first time that favipiravir inhibited JUNV through the accumulation of transition mutations, confirming its role as a purine analogue against arenaviruses. Moreover, we showed that treatment with a combination of favipiravir and either ribavirin or remdesivir inhibited JUNV replication in a synergistic manner, blocking the generation of the drug-resistant mutant. Our findings provide new insights for the clinical management and treatment of Argentine hemorrhagic fever.

## INTRODUCTION

Argentine hemorrhagic fever (AHF) is a severe zoonotic disease caused by Junin virus (JUNV) and highly endemic in Argentina. In addition to the intense clinical course of the disease, the lack of approved therapeutics and preventive countermeasures against JUNV highlight its significant threat to global public health (NIAID Emerging Infectious Diseases/Pathogens; Borio et al., 2002; Enria et al., 2008). There are a few reports on an immune plasma therapy and a combinational, off-label use of ribavirin and favipiravir (Enria et al., 2008; Veliziotis et al., 2020). Although a live attenuated vaccine, Candid #1, was developed by the US Army Medical Research Institute of Infectious Diseases, it has been approved only for use in endemic areas due to concerns over its genomic stability (Gowen et al., 2021; McKee et al., 1993; Stephan et al., 2013).

Favipiravir (6-fluoro-3-hydroxy-2-pyrazinecarboxamide; also known as T-705 or Avigan) is a purine analogue originally developed as an antiviral agent for influenza and subsequently reported to inhibit the replication of a broad spectrum of RNA viruses (Delang et al., 2018). Favipiravir is a prodrug that is metabolized into its active form, ribofuranosyl 5′-triphosphate (favipiravir-RTP), upon cellular uptake, and thus acts as a pseudo-nucleotide that competes with endogenous guanine and adenine nucleotides, leading to the disruption of viral replication and transcription (Furuta et al., 2005; Goldhill et al., 2019). Given the extensive structural and functional similarities among RNA-dependent RNA polymerase (RdRp) of RNA viruses (Bruenn, 2003), favipiravir remains a promising countermeasure against emerging and re-emerging viral diseases caused by RNA viruses. Clinical trials showed the efficacy of favipiravir against viral hemorrhagic fever caused by the Severe Fever with Thrombocytopenia Syndrome Virus (SFTSV), and a direct correlation between favipiravir treatment and reduction in viral RNA levels (Suemori et al., 2021). However, an open-label observational study on Ebola virus showed lack of efficacy of favipiravir treatment in reducing the viral load reduction and improving survival of the affected subjects (Madelain et al., 2017). Prior to the discovery of favipiravir, ribavirin was the only available antiviral drug effective against JUNV (Weissenbacher et al., 1986). However, several concerns over safety and efficacy are linked with the use of ribavirin, limiting its clinical use (Enria et al., 2008). Several pre-clinical studies investigated the inhibitory effect of antiviral drugs in JUNV infections (Gowen et al., 2017, 2013). Favipiravir showed high protection against lethal JUNV infection and was well tolerated at high doses. Notably, while ribavirin shows high efficacy in suppressing viral replication, the mortality of the infected animal models is only delayed or slightly reduced by ribavirin (Kenyon et al., 1986; McKee et al., 1988). Moreover, while favipiravir specifically targets the viral polymerase with minimal side effects (Furuta et al., 2005), ribavirin acts through multiple mechanisms. In addition to the inhibition of viral polymerase, ribavirin targets cellular inosine monophosphate dehydrogenases and restricts intracellular GTP availability, thereby indirectly inhibiting virus replication. This explains the synergistic effect of ribavirin when used with favipiravir, which offers a combinational therapeutic approach for the clinical management of patients with AHF (Carrillo-Bustamante et al., 2017; Westover et al., 2016).

The emergence of drug-resistant mutants invalidates the effect of antiviral drugs in the short term. To date, experimental isolation of favipiravir-resistant mutants has only been reported for chikungunya virus (Delang et al., 2014) and enterovirus 71 (Wang et al., 2016), which are positive-sense RNA viruses, as well as the influenza A virus, which contains a negative-sense, segmented RNA genome (Goldhill et al., 2018b). Isolation of drug-resistant mutants represents a useful approach for studying the molecular mechanisms of antiviral drugs (Lo et al., 2020). Considering the possibility of the emergence of drug-resistant mutants and the gap in the mechanistic data on the antiviral action of favipiravir against arenaviruses (Mendenhall et al., 2011b), we attempted to isolate favipiravir-resistant JUNV. In this study, we isolated favipiravir-resistant mutants, also labelled escape mutants, with two amino acid substitutions on the viral GPC and RdRp. The analyses of the escape mutant, for the first time, provided experimental evidence that favipiravir primarily acts against JUNV by inducing transition mutations. Here, we showed that the selective pressure of favipiravir promoted the emergence of the JUNV variant with a higher replication fidelity and, therefore, a lower susceptibility to favipiravir. We also showed that the treatment with combination of favipiravir with either ribavirin or remdesivir inhibited JUNV replication in a synergistic manner. The clinical implications of these findings need to be considered prior to the therapeutic use of favipiravir in the individuals with a recent history of Candid #1 vaccination.

## RESULTS

### Isolation of favipiravir-resistant mutants

For the isolation of favipiravir-resistant JUNV mutants in this study, 293T cells were chosen because of the robust replication kinetics of JUNV in this cell line. To identify the optimal selective pressure of favipiravir on JUNV replication, the IC_50_ value was determined by a dose-response experiment. 293T cells were infected with Candid #1 virus at a multiplicity of infection (MOI) of 0.1, in the presence of a series of favipiravir dilutions or DMSO. Quantification of viral titers at 48 hours post infection (hpi), showed favipiravir IC_50_ to be 4.9 µM with 95% confidence interval (CI) of 4.5 µM to 5.4 µM. This value is comparable with that of a previous study reporting favipiravir IC_50_ for JUNV in Vero cells (Gowen et al., 2007). The cytotoxicity assay showed that favipiravir lacked toxic effects on cell viability at the specified concentrations (**Fig. S1**). We selected 5 µM of favipiravir as an optimal selective pressure. First, the cells were infected with the JUNV Candid #1 strain (MOI: 0.01). After adsorption, medium containing either favipiravir or DMSO was added, and cells were then incubated. Additional infection control using fresh virus stock (no passage control) in the absence of favipiravir/DMSO was used to monitor the accumulation of defective interfering particles (data not shown) (Ziegler and Botten, 2020). Viral titers were measured using plaque assays after every passage. On the first two initial passages, P0 and P1, JUNV titers showed an 81.7- and 105-fold reduction, respectively (**Fig. 1A**). To increase the selective pressure, favipiravir concentration was increased to 20 µM (approximately IC_90_) for subsequent passages. The increase of favipiravir concentration was associated with a reduction in titer similar to that detected using 5-µM favipiravir, indicating that a resistant population was beginning to emerge. As shown in **Fig. 1A**, at passage 11 (P11), viral titers were similar to those of no-drug controls, suggesting that a dominant proportion of the viral population was resistant to favipiravir. Further passaging (P12 and P13) in the presence of favipiravir did not affect viral titers. To measure the reduction in susceptibility of the P11 virus to favipiravir, a dose-response assay was performed. As shown in **Fig. 1B**, IC_50_ values of P0 and P11 were 4.3 µM (95% CI = 3.8 to 4.9) and 27.1 µM (95% CI = 23.1 to 31.6), respectively, indicating a significant increase in IC_50_ value (6.3-fold) and the emergence of favipiravir-resistant Candid #1-mutant (Candid #1-res).

**Figure 1.**
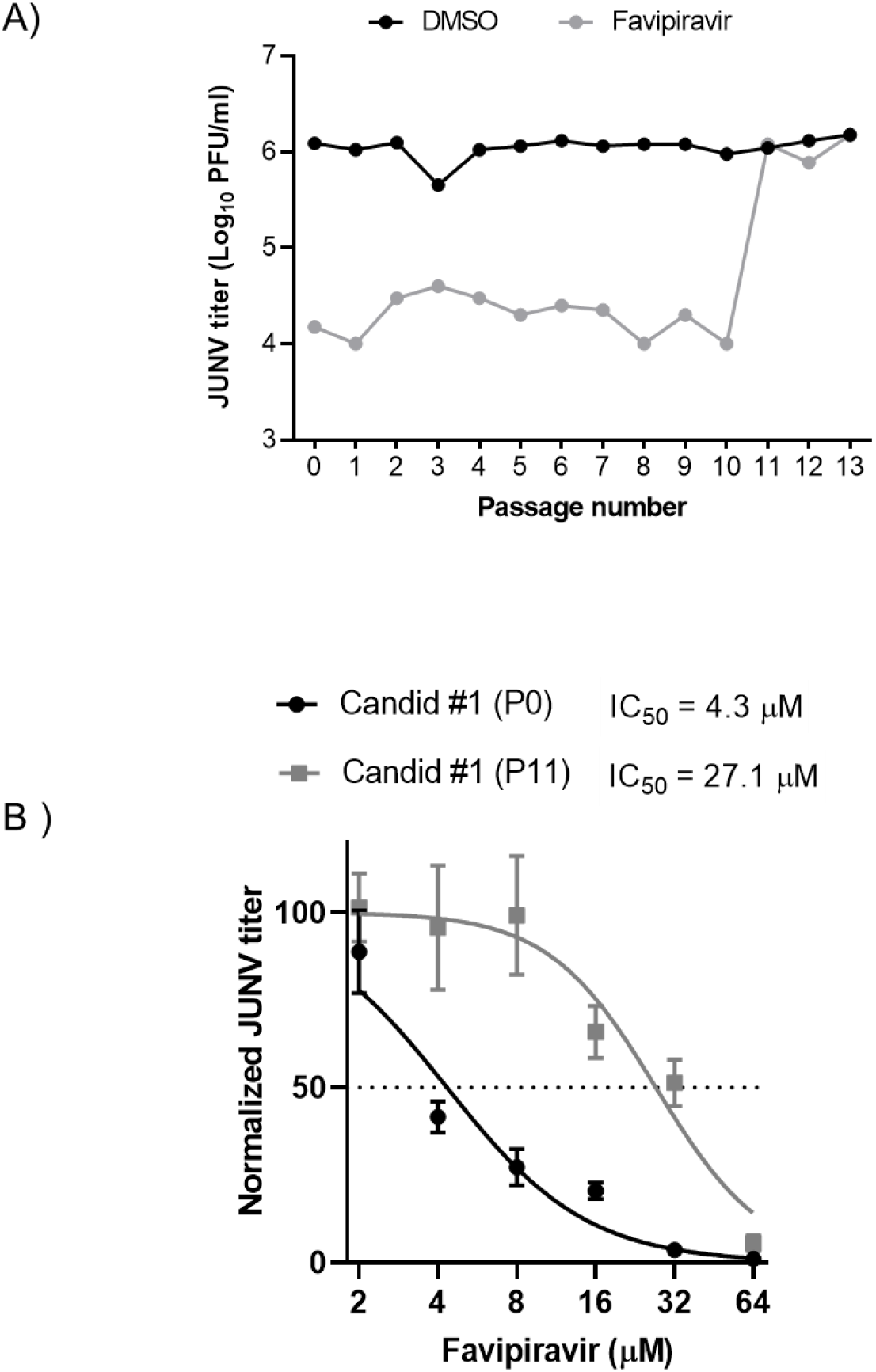
Emergence of favipiravir resistant JUNV by sequential passaging in 293T cells. (A) Serial passage of JUNV in presence of moderate favipiravir concentrations. JUNV was passaged in 293T cells (MOI: 0.01) in presence of 5 µM for the first three passages and 20 µM favipiravir for the remaining passages. After a total of 11 passages, a resistant JUNV population emerged (gray). Control passages were performed in parallel using DMSO (black), n = 1. (B) Favipiravir dose-response analysis. 293T cell were infected with JUNV Candid #1 parental (P0) or passage 11 (P11) viral populations (MOI: 0.1), After adsorption, media containing indicated concentrations of favipiravir was added to the cells. At 48 hpi., viral titers were measured by plaque assay. Error bars indicate ±SD; three independent experiments in duplicate (n = 6) were performed; nonlinear regression analysis was applied; LOD, limit of detection.

### Identification of the mutations in the favipiravir-resistant JUNV mutant

To identify the mutations that confer resistance to favipiravir, four clones were isolated from P11 by plaque assay, as described in the Materials and Methods section. Viral RNA from each clone was extracted and the sequences of all open reading frames (ORFs) were determined. All four clones showed the same mutations as the P0 parental virus. Three nucleotide substitutions, two in the RdRp coding region (1384 A to G, 3669 A to G) and one in the GPC coding region (502 G to A), were identified in P11 mutants **(Fig. 2)**. In the RdRp region, the A to G substitution at 3669 generated a synonymous mutation, whereas the A to G substitution at 1384 led to N462D amino acid modification in the PA-like domain of RdRp (Brunotte et al., 2011; Peng et al., 2020). In the GPC region, G to A substitution at 502 caused anA168T amino acid substitution within the GP1 subunit. No mutations were observed in Z or NP genes.

**Figure 2.**
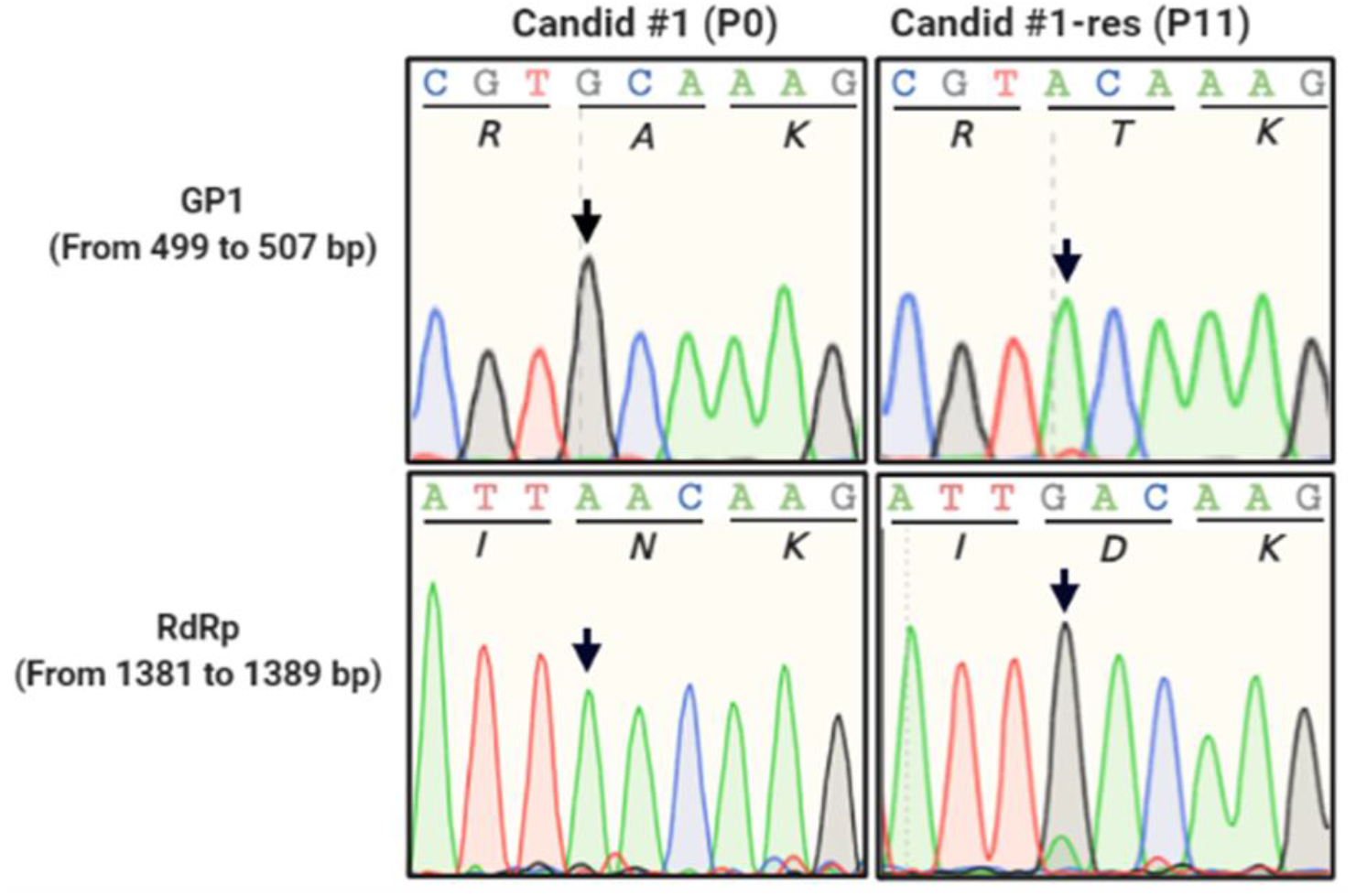
Nucleic acid substitutions in RdRp and GPC open reading frames. Representative chromatograms of mutations of favipiravir resistant JUNV Candid #1, passage 11 (P11) are shown in comparison to the parental virus population (P0). Amino acid residues are shown below each codon. Arrows indicate mutation locations.

### Growth kinetics of favipiravir-resistant JUNV mutant

First, we compared the growth kinetics of Candid #1-res (P11) to parental Candid #1 (P0) in the absence of favipiravir. After virus adsorption on ice to synchronize the infection as described in the Materials and Methods, 293T cells infected with either Candid #1 or Candid #1-res were incubated and culture supernatants were collected at 8, 12, 24, and 28 hpi Viral titers in the culture supernatants were determined by plaque assay. Efficient replication of Candid #1-res was observed at 8 hpi, which was much earlier than that of parental Candid #1 (**Fig. 3**). However, there was no significant difference between the titers of both viruses at 28 hpi, indicating that Candid #1-res exhibited the rapid growth. No differences in plaque morphologies were observed (data not shown).

**Figure 3.**
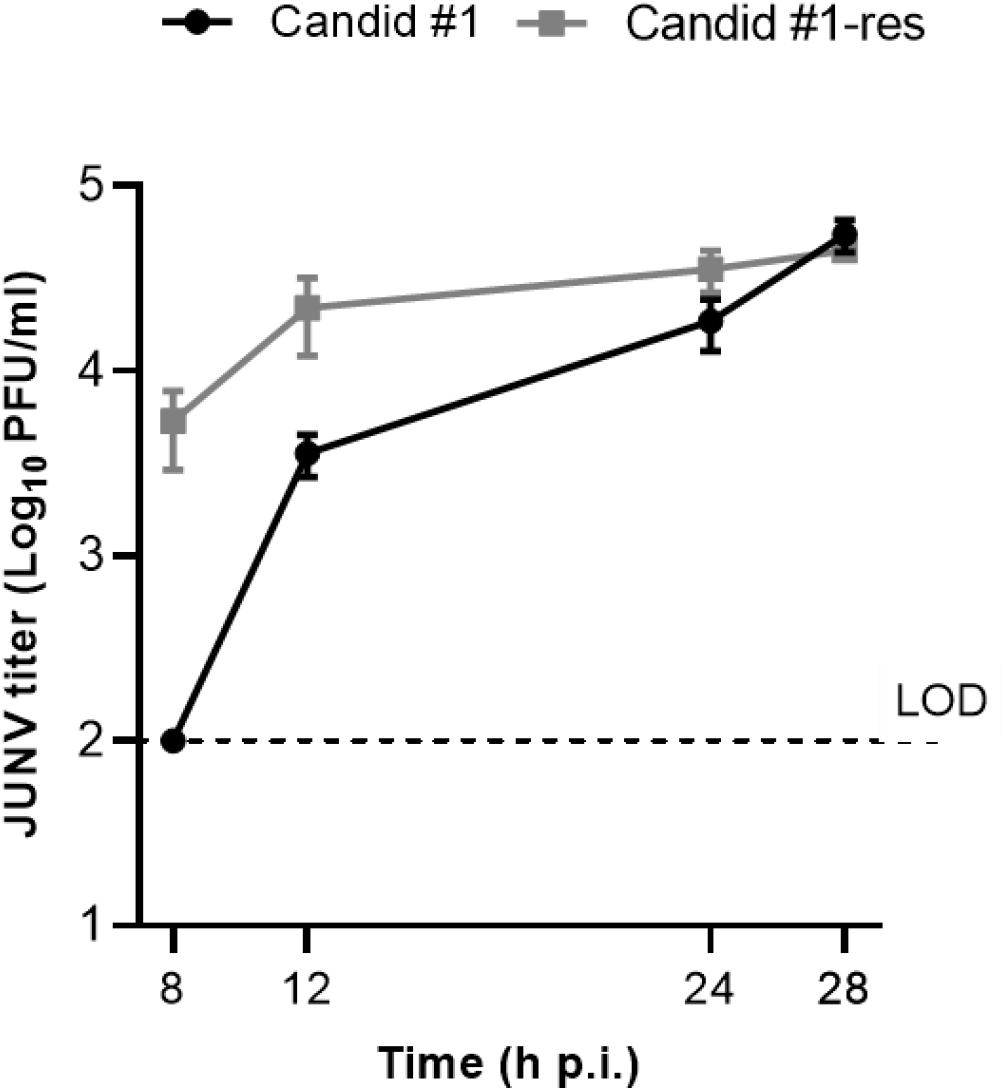
JUNV Candid #1-mutant virus replication begins at earlier time point. To determine the one-step growth kinetics of favipiravir resistant JUNV, 293T cell were infected with either Candid #1 or Candid #1-mutant (MOI: 0.1). In order to synchronize infection, cells were incubated on ice for 30 minutes. Cells were then washed with PBS (−) and pre-warmed DMEM containing 10% FBS was added. Supernatant was collected at the indicated time points. Titers were determined by plaque assay. Error bars indicate ±SD, three independent experiments in duplicates (n = 6) were performed. Statistical significance was determined by 2-way ANOVA test (*P* < 0.0001).

### High-fidelity replication of favipiravir-resistant JUNV

To understand the mechanism by which JUNV acquired resistance against favipiravir, the mutation frequency of Candid #1 and Candid #1-res in the presence of favipiravir was assessed. First, 293T cells were infected with Candid #1 or Candid #1-res at MOI=0.1 and incubated in the presence of 20 µM favipiravir. After 48 h, the culture supernatants were collected. The nucleoprotein (NP) gene of arenaviruses is highly prone to mutations (Grande-Pérez et al., 2015), therefore, we selected and cloned the same region of the NP gene from each virus and performed clonal sequencing. As shown in **Fig. 4**, the mutational analysis of a total of 24,300 nucleotides for the parental Candid #1 and 27,450 nucleotides for the Candid #1-res showed that the resistant mutant had acquired 1.82 mutations per 10,000 nucleotides as compared to the 8.64 mutations of the parental population (4.7-fold lower. *P* = 0.0053, Mann– Whitney *U* test), indicating higher fidelity of the Candid #1-res virus. Similarly, mutation frequencies of Candid #1-res in DMSO-treated controls were slightly lower (3.8-fold), although the data lacked statistical significance (*P* = 0.242, Mann–Whitney *U* test). The frequency of favipiravir-induced mutations in the parental Candid #1 is estimated to be beyond the tolerable threshold of error-catastrophe, and is comparable to those reported for other viruses, including influenza, Zika, and murine norovirus treated with favipiravir (Arias et al., 2014; Bassi et al., 2018; Goldhill et al., 2019). These data suggest that mutagenesis with lethal consequences is the primary mechanism of favipiravir antiviral action against JUNV.

**Figure 4.**
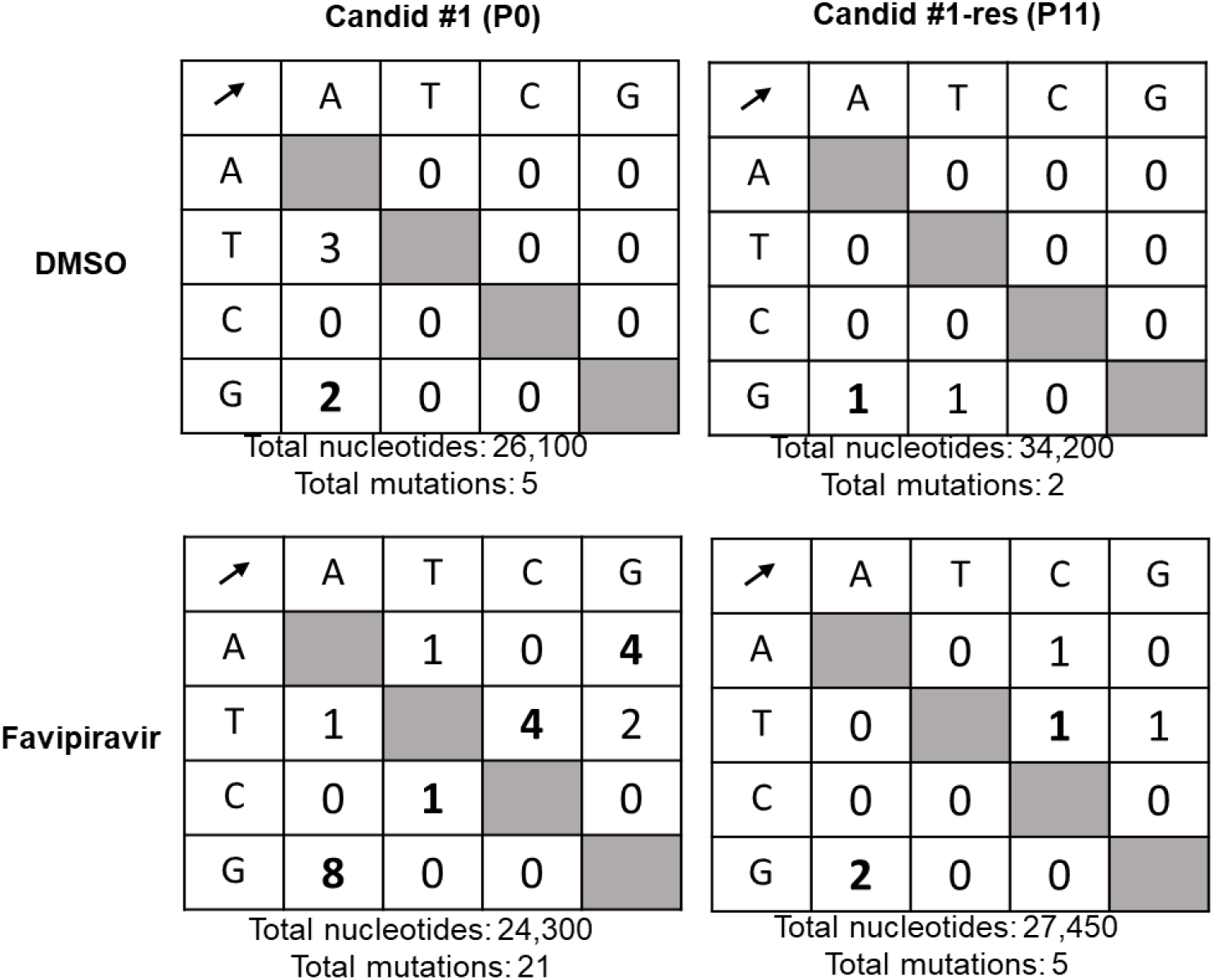
Mutational frequencies of virus populations in presence of 20 µM favipiravir or DMSO control. Clonal sequencing targeting 450 bp of nucleoprotein (NP) gene was performed (as described in materials and methods) to estimate the frequency of mutations in each virion population. Nucleotide polymorphisms were counted for Candid #1 and Candid #1-res viruses to estimate the proportions of transition (shown in bold) and transversion substitutions in presence of 20 µM favipiravir.

Next we categorized the substitutions to determine the proportions of transitions versus transversions. The most common mutations in the presence of favipiravir were G to A, followed by T and C mutations, accounting for 80% of all substitutions in favipiravir-treated viruses, indicating that the mutation profile induced by favipiravir is biased towards transitions **(Fig. 4)**. This is in agreement with studies on other viruses treated with favipiravir (Arias et al., 2014; Ávila et al., 2016; Goldhill et al., 2019; Guedj et al., 2018). Consistent with the literature, supplementation of purines, but not pyrimidines, reversed the antiviral activity of favipiravir **(Fig. S2)** (Mendenhall et al., 2011a), reaffirming the role of favipiravir as a purine analogue that competes with adenosine and guanosine during nucleotide incorporation. This, in turn, explains the error bias observed in the transitional mutations.

### Reduced specific infectivity of favipiravir-resistant JUNV

To further evaluate the mutagenic effect of favipiravir on Candid #1 and Candid #1-res viruses, their respective specific infectivity (defined as the ratio of infectious virions to the encapsidated genome copy number), when exposed to increasing concentrations of favipiravir, were compared. 293T cells were infected with Candid #1 or Candid #1-res (MOI=0.01), and then treated with serial dilutions of favipiravir or DMSO. Specific infectivity of Candid #1 at 48 hpi was 0.85 and 0.39 log_10_ plaque-forming units (PFU) per mL/log_10_ RNA copies per mL for 2 µM and 64 µM favipiravir, respectively, and showed a concentration-dependent reduction **(Fig. 5**). This suggests that increasing concentrations of favipiravir cause the accumulation of lethal mutations that lead to the loss of JUNV infectivity (Arias et al., 2014; Espy et al., 2019). In contrast, specific infectivity of Candid #1-res virus was only slightly affected from 1.009 to 0.80 log_10_ PFU per mL/log_10_ RNA copies per mL, when exposed to 2-µM and 64-µM favipiravir respectively. The significant reduction in specific infectivity of Candid #1 compared to that of the Candid #1-res virus (*P* < 0.0001, two-way ANOVA test) suggests that the resistant mutant is less susceptible to the mutagenic effect of favipiravir. Notably, we did not observe any difference in viral RNA copy numbers between Candid #1 and Candid #1-res variants across any concentration (up to 64 µM) of favipiravir treatment (data not shown). Taken together, these data indicate that the lower susceptibility of Candid #1-res virus to favipiravir is mediated by its higher replication fidelity.

**Figure 5.**
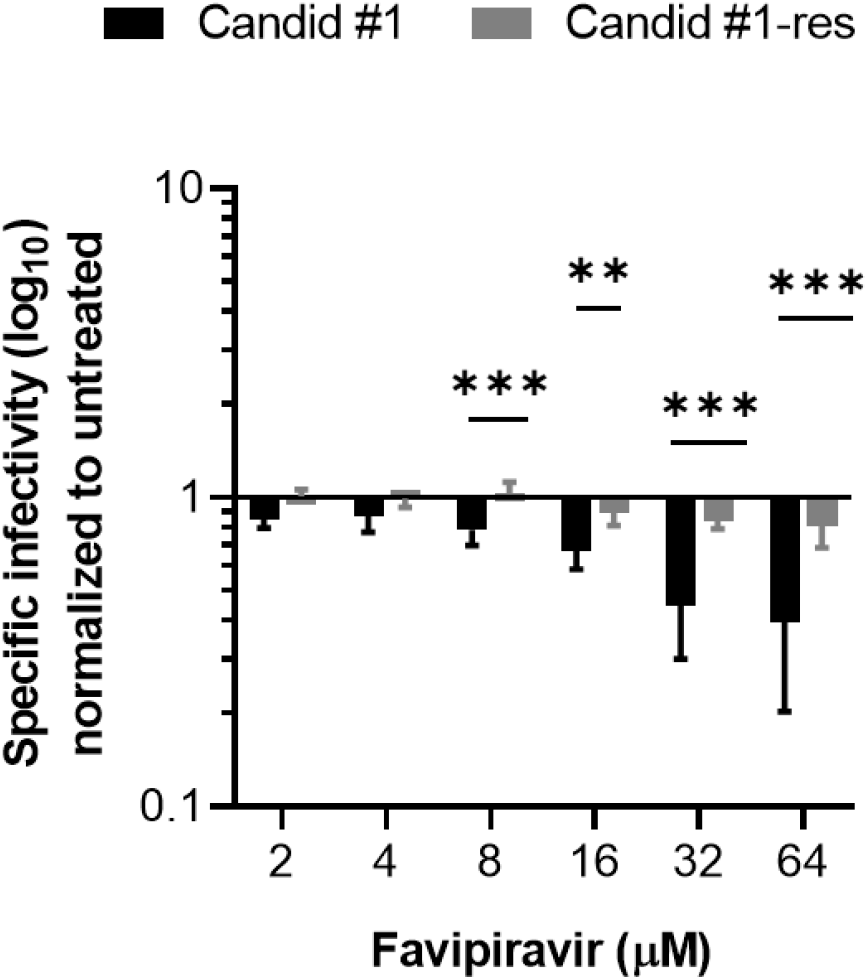
Infectivity of Candid #1 and Candid #1-res viruses. 293T cells were infected with each virus (MOI: 0.01) and treated with different concentrations of favipiravir or DMSO **S**pecific infectivity values at 48 hpi, were calculated using the ratio of infectious particles (Log_10_ PFU/mL) to the RNA copy numbers (Log_10_ copies/mL). Values were normalized to DMSO-treated controls. Error bars indicate ±SD; two independent experiments in duplicates (n = 8) were performed. Statistical significance was determined by 2-way ANOVA test (** indicates *P* < 0.01 and *** indicates *P* < 0.001).

### Enhancement of JUNV growth by GPC-A168T substitution

To investigate the functional impact of GPC A168T substitution on virus entry, we compared the internalization dynamics of favipiravir-susceptible (P0) and resistant variants (P11) using a pseudotyped vesicular stomatitis virus (VSV) system in 293T cells. To normalize the number of viral particles used for infection, real time qPCR targeting the VSV-M gene was performed **(Fig. S3)**. A confluent monolayer of cells was infected with the pseudotyped VSV bearing GPC-A168 or GPC-T168, Candid#1pv-A168 or Candid#1pv-T168, which have equivalent copies of the VSV genome. To allow synchronized virus entry, cells were first incubated at 4 °C for 30 min and subsequently transferred to 37 °C for further incubation. Measurements of luciferase signal at 8, 16, and 24 hpi showed a significant difference at 16 hpi, with Candid#1pv-T168 having more robust entry kinetics compared to Candid#1pv-A168 **(Fig. 6A)**. We then examined the intracellular levels of the VSV-M protein as a marker of fusion efficiency. Using a similar experimental setup, 293T cells were infected with an equal number of viral particles (input virus). After 8 h, the cells were lysed, and samples were prepared for the detection of VSV-M protein using western blotting **(Fig. 6B)**. Candid#1pv-T168 showed 12 times more intracellular M protein expression levels than Candid#1pv-A168, despite similar levels of input virus **(Fig. 6C)**, indicating that the viral genome was released into the cytoplasm more efficiently, leading to more rapid and elevated VSV-M protein expression. These data suggest that Candid#1pv-T168 is more efficient in the entry and/or fusion processes, leading to an altered viral life cycle.

**Figure 6.**
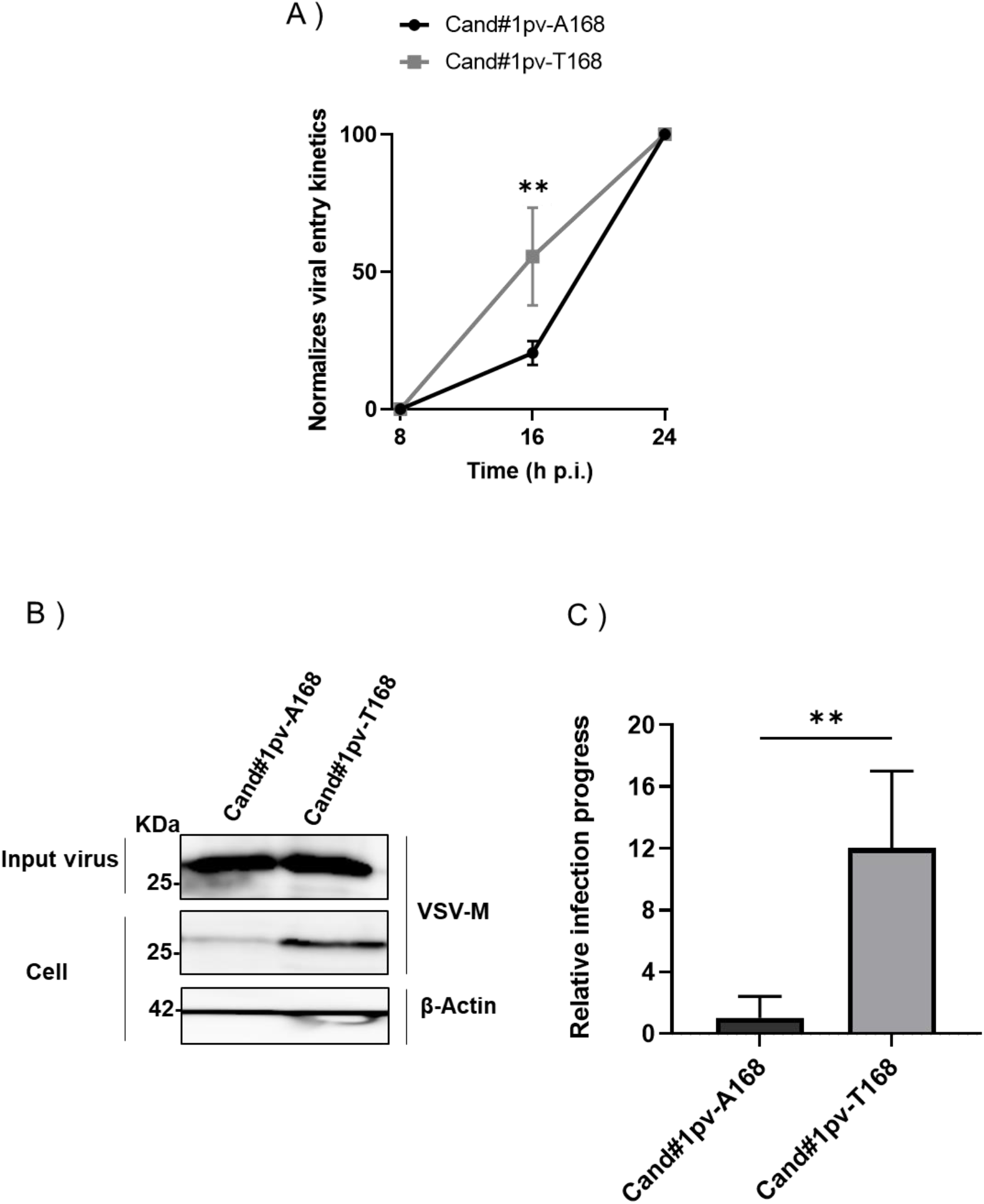
GPC-A168T substitution enhances viral entry dynamics. (A) 293T cells were infected with Candid#1pv-A168 or Candid#1pv-T168 virus at 4 °C to allow synchronized attachment. Unbound viral particles were removed and internalization was initiated by transferring the cells to 37 °C. Luciferase signal was measured at indicated time points. Signal at 24 hours was considered 100%. (B) Western blot analysis (anti-VSV-M in the upper panel for the detection of the VSV-M protein, and Anti-β-Actin as loading control) to assess fusion efficiency of pseudotyped viruses (C) Expression levels of intracellular M protein was normalized to the input virus. Quantified results of two independent experiments (n = 6) are shown. The bar indicates ±SD. Statistical significance was determined using multiple *t*-tests (** indicates *P* < 0.01).

### Effect of RdRp-N462D substitution on RNA polymerase activity

Next, we investigated whether N462D substitution affects RdRp activity in Candid #1. A minigenome (MG) system based on the S segment of Candid #1 virus was constructed, and an MG assay in absence of favipiravir was performed. To ensure that both RdRp-N462 and - D462 plasmids had comparable expression levels, both proteins were tagged with a FLAG peptide and similar expression levels were confirmed by western blot analysis **(Fig. 7A** and **B)**. 293T cells were transfected in 24-well plates with the plasmids for either NP, MG, RdRp-N462, RdRp-D462, or empty vector. Luciferase signals (normalized to the internal control) were measured at 24 hours post-transfection (hpt). The results are expressed as relative induction rates. We observed that N462D substitution had no significant effect on RNA polymerase activity **(Fig. 7C)**. To examine the sensitivity of RdRp-D462 to favipiravir, the activity of both polymerases exposed to different concentrations of the drug using a minigenome system was tested. No significant difference in favipiravir dose-response in RdRp-N462 (IC_50_ = 245.6–304.3 µM [95% CI]) and RdRp-D462 (IC_50_ = 289.8–362.1 µM [95% CI]) was observed, although RdRp-D462 showed a slightly increased resistance to favipiravir compared with that of RdRp-N462 **(Fig. 7D)**. No cytotoxicity was observed in this assay **(Fig. 7E)**. Quantification of the luciferase mRNA showed that high concentrations of favipiravir (≥200 µM) act as a chain terminator, and reflected the reduction of luciferase signal in the MG assay **(Fig. 7F)**.

**Figure 7.**
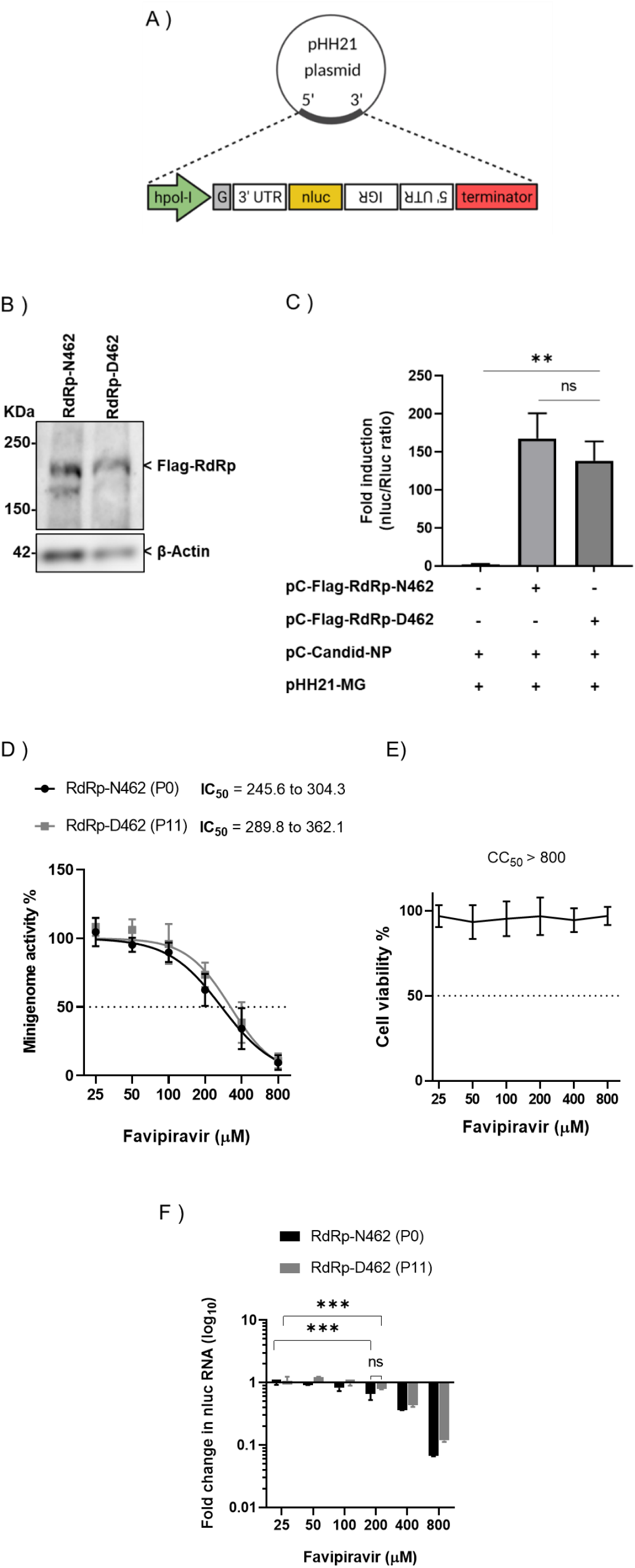
Effect of RdRp-N462D substitution on polymerase activity. (A) Schematic representation of the minigenome (MG) plasmid constructs with a nluc reporter (B) Western blot analysis (anti-Flag; upper panel for the detection RdRp-N462 or RdRp-D462 protein. Anti-β-Actin; loading control) to ensure equal expression levels (C) Polymerase activities measured using the MG system in 293T cells at 48 hpt. Results are expressed by the ratio of nano luciferase to renilla luciferase activity (internal control). Quantified results of two independent experiments (n = 6) are shown. The bar indicates ±SD. Statistical significance was determined using *t*-tests (** indicates *P* < 0.01. ns indicates not significant). (D) Sensitivity of RdRp-N462 and RdRp-D462 polymerases were compared using the MG system. Quantified results of two independent experiments (n = 6) are shown. (E) Cell viability assay as described in materials and methods. (F) Quantification of nLuc mRNA from a minigenome assay by qPCR. ΔΔCT was calculated using *GAPDH* as described in the materials and methods. Fold change in mRNA was normalized to DMSO-treated controls. Error bars indicate ±SD; two independent experiments in duplicates (n = 6) were performed. Statistical significance was determined using a 2-way ANOVA test (*** indicates *P* < 0.001).

### Combinational inhibitory effect of favipiravir with either ribavirin or remdesivir on JUNV growth

Combination antiviral therapy is a promising approach to minimize the risk of the emergence of drug resistance and enhance the antiviral effect. Therefore, we investigated the inhibitory effect of favipiravir in combination with ribavirin and remdesivir. As shown in **Fig. 8**, the anti-JUNV effect of favipiravir was significantly higher when combined with ribavirin (ZIP synergy score: 14.02) or remdesivir (ZIP synergy score: 15.82) without any significant antagonistic effect. No cytotoxicity was associated with any of the tested drug combinations. Despite our attempt to isolate a resistant variant to combinational treatments, no resistant variant was generated even after 15 passages **(Fig. S4)**.

**Figure 8.**
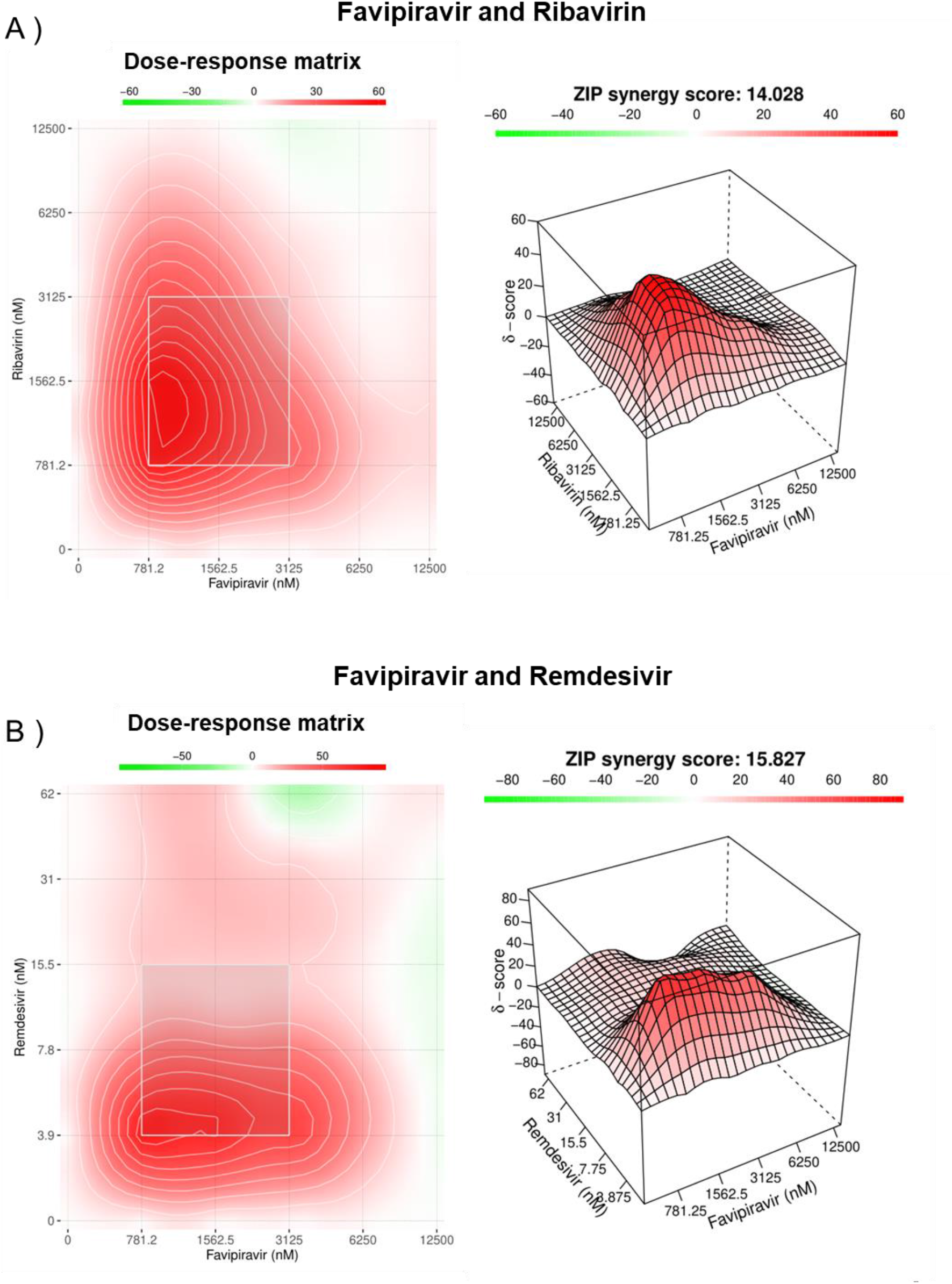
Combination inhibitory effect of favipiravir and ribavirin or remdesivir on JUNV. 293T were infected with JUNV (MOI, 0.1) and subsequently treated with a 6 × 6 drug combination matrix of favipiravir + ribavirin (A) or favipiravir + remdesivir (B) Dose-response matrix and synergy heat map are presented. Colored bar indicates strength of synergy (δ-score); less than −10 is likely to be antagonistic, −10 to 10 suggests an additive drug interaction, larger than 10 indicates a synergistic effect. Data are means of two independent experiments in duplicates (n = 4).

## DISCUSSION

In this study, we attempted to understand the mechanism of antiviral action of favipiravir against JUNV by isolating resistant variants. In our approach, lower concentrations of favipiravir for three initial passages followed by higher concentrations for the remaining passages were used. This allowed a gradual accumulation of mutant variants under a moderate drug pressure and to avoid sudden exposure of JUNV to lethal concentrations of favipiravir (Pauly and Lauring, 2015), thus enabling us to successfully maintain and isolate the resistant population.

The arenavirus RdRp consists of three domains: an N-terminal PA-like domain with endonuclease activity, a polymerase region possessing the active site, and a PB2-like domain. In contrast to previous studies showing that RNA viruses developed resistance to favipiravir through mutations in the conserved catalytic domain of viral RdRp (Delang et al., 2014; Goldhill et al., 2018b; Wang et al., 2016), we identified an RdRp-N462D substitution within the PA-like domain in favipiravir-resistant variants. A recent study, which resolved the structure of arenavirus polymerase protein with a near-atomic resolution, showed that residue 462 of new world arenaviruses belongs to the core lobe region of the PA-like domain, which is involved in stabilization of the polymerase active site. However, the precise interactions of the =462 residue are yet to be clarified (Peng et al., 2020). Our assessment of the functional impact of N462D substitution using an MG system showed only a slight, statistically non-significant (*P* = 0.125) reduction in reporter activity without any major impact on polymerase function **(Fig. 7C)**. However, comparisons of mutation frequencies of Candid #1 and Candid #1-res virus revealed a significant reduction in polymerase error number in the case of RdRp-D642, suggesting an important role of this residue in polymerase fidelity (**Fig. 4)**. The leading hypothesis on the mechanism of the favipiravir resistance observed in this study is that the N462D substitution enhances the selectivity of RdRp for the correct nucleoside triphosphates during replication and transcription, resulting in lower favipiravir incorporation, as described for other mutagen-resistant RNA viruses (Cheung et al., 2014; Pfeiffer and Kirkegaard, 2003). Further analysis of binding affinities will clarify the precise mechanism of D462 resistance to favipiravir. Notably, the higher fidelity of Candid #1 mutant virus correlated with resistance to favipiravir, and the virus remained susceptible to higher concentrations of the drug, indicating that RdRp-D462 does not tolerate the chain termination activity of favipiravir, as was demonstrated in this study through the MG system **(Fig. 7D and F)**. Accordingly, the Candid #1-res virus remained susceptible to other purine analogues, ribavirin and remdesivir, with non-mutagenic mechanisms of action **(Fig. S5)** (Feld and Hoofnagle, 2005; Furuta et al., 2005; Mendenhall et al., 2011a; Tchesnokov et al., 2020). To date, with the exception of the influenza virus (Cheung et al., 2014), other RNA viruses with high replication fidelity possess a positive-sense, non-segmented genome (Pfeiffer and Kirkegaard, 2003; Sadeghipour et al., 2013). To the best of our knowledge, this is the first report on the isolation of a high-replication fidelity phenotype amongst hemorrhagic fever viruses. Studies have demonstrated that higher fidelity of replication affects the genetic heterogeneity of viral sub-populations, imposing a fitness cost *in vivo* (Cheung et al., 2014; Pfeiffer and Kirkegaard, 2005; Vignuzzi et al., 2006). Hence, there remains a need to further investigate the virulence and pathological characteristics of JUNV with high-fidelity replication, which was isolated in this study.

The other mutation (A168T) identified in this study was found to be within the GP1 subunit of the glycoprotein complex (GPC). Arenavirus GPC is a precursor protein that forms a trimer of stable signal peptides, GP1 and GP2 subunits, upon maturation by cellular enzymes. During virus entry, GP1 and GP2 are responsible for the recognition of receptors and the fusion with endosome membranes, respectively (Urata and Yasuda, 2012). While A168T substitution led to more efficient viral entry **(Fig. 6)**, no impact on attachment of pseudotype viral particles to the target cells was observed (data not shown), suggesting that the functional importance of the A168T substitution is on the post-attachment step of JUNV virus entry. Consistent with this, growth kinetics of the Candid #1-res virus represented more robust replication at earlier time points **(Fig. 3)**. In recent years, a novel mechanism of drug resistance mediated by an altered viral life cycle has been postulated (Neagu et al., 2018; Sedaghat and Wilke, 2011). While there is no experimental evidence to fully support this theory, co-emergence of surface glycoprotein mutations together with RdRp mutation also has been reported to occur in a remdesivir-resistant variant of SARS-CoV-2 (Szemiel et al., 2021), highlighting the possible role of infection synchronicity (life cycle adaptability) on the potency of antiviral drugs. Nevertheless, in the absence of a reverse genetics system, we were unable to confirm whether the altered life cycle of JUNV imposed by the GP1-A168T substitution plays a direct role in reducing susceptibility to favipiravir.

Here, we experimentally demonstrated that the potency of favipiravir could be significantly enhanced against JUNV if used in combination with ribavirin or remdesivir **(Fig. 8)**. Furthermore, we showed that it was difficult to isolate JUNV variants that were resistant to the combination treatment **(Fig. S4)**. These findings suggest the potential of combination therapies for favipiravir with ribavirin or remdesivir.

In conclusion, we described the isolation of a high replication fidelity variant of arenavirus with reduced susceptibility to favipiravir. More importantly, we provide experimental evidence that hyper-mutagenesis is the primary mechanism of favipiravir action against JUNV. Consistent with our observations, studies on favipiravir treatment of non-human primates infected with Lassa virus showed a reduction in virus infectivity without affecting viral load, providing evidence that favipiravir is primarily a mutagen against old world arenaviruses (Lingas et al., 2021; Rosenke et al., 2018). Our findings emphasize the importance of the addition of a non-mutagenic inhibitor to the treatment regimens for the Argentine hemorrhagic fever (AHF).

## MATERIALS AND METHODS

### Cells, viruses, and compounds

Human embryonic kidney (293T) and African green monkey kidney (Vero 76) cell lines were maintained in Dulbecco’s modified Eagle’s medium (DMEM; Invitrogen, CA, USA) with 10% fetal bovine serum (FBS) and 1% penicillin and streptomycin. The Candid #1 vaccine strain of JUNV was kindly provided by Dr. Juan C. de la Torre (Scripps Research Institute, California, USA). Favipiravir was obtained from FUJIFILM Toyama Chemical CO., LTD. (Toyama, Japan). Ribavirin (Sigma Aldrich, MO, USA) and remdesivir (Cayman, MI, USA) were purchased. All compounds were dissolved in 100% dimethyl sulfoxide (DMSO) and stored at −30 °C until use.

### Virus infection and titration

Cells were infected with Candid #1 strain at the indicated multiplicity of infection (MOI). After adsorption for 1 h at 37 ?, the inoculum was removed and washed with PBS (−). Pre-warmed DMEM containing 10% FBS was added to the cells, which were then incubated at 37 °C with 5% CO_2_. For the quantification of viral titers, a plaque assay was performed according to standard procedures using 10-fold dilutions of the samples in Vero 76 cells as previously described (Zadeh et al., 2020).

### Determination of inhibitory concentrations and toxicity testing

To determine the half-maximal inhibitory concentration (IC_50_), 293T cells were infected at a MOI of 0.1 in 24-well plates as explained above. After adsorption, the virus solutions were removed, and fresh DMEM containing serial dilutions of compounds (ranging from 2 µM to 64 µM for favipiravir/ribavirin and 0.0125 µM to 4 µM for remdesivir) were added to the infected cells. At 48 hpi, the supernatants were collected to determine viral titers by plaque assay. To plot the dose-response curve, viral titers from each drug concentration were normalized to the titers in the DMSO control. The cytotoxicity of the compounds was assessed using the CellTiter-Glo cell viability assay (Promega, Madison, WI, USA), following the manufacturer’s instructions. Briefly, 293T cells were seeded in a 96-well plate and incubated overnight. Cells were then treated with different concentrations of each compound, as described above. After 48 h, CellTiter-Glo reagent was added, and luminescence was measured using an illuminometer (Tristar LB941, BERTHOLD). Cell viability in DMSO-treated controls was set to 100%.

### Selection and purification of JUNV favipiravir-resistant mutants

To isolate favipiravir-resistant JUNV, we serially passaged the Candid #1 strain in 293T cells at a MOI of 0.01 under the selective pressure of favipiravir (5 µM for the first three passages and 20 µM for the remaining passages). As a control, viruses were serially passaged in the absence of favipiravir in parallel. Supernatants were diluted 10 times in Opti-MEM (Invitrogen) before infecting the cells for the next passages. At 48 hpi, two aliquots of the supernatants were prepared and stored at −80 ?. Virus titers were measured using plaque assays. To isolate a single clone of the virus, a plaque assay was performed in 6-well plates in Vero 76 cells as described above. After 7 days of incubation, plaques were collected and inoculated into 293T cells to expand the virus clone. To isolate resistant mutants against combination of favipiravir (0.3 µM) and ribavirin (0.3 µM) or remdesivir (1 nM), the virus was passaged and titrated under similar conditions as stated above.

### Reverse transcription polymerase chain reaction (RT-PCR) and RNA sequencing

RNA was extracted from the supernatant of cells infected with Candid #1 (P0 and P11) using the QIAamp Viral RNA Mini Kit (Qiagen, Hilden, Germany), according to the manufacturer’s instructions. For the sequencing, 15 sets of primers were designed to produce overlapping PCR products of 800 to 900 bp (Table S1) using the Primal Scheme (available at http://primal.zibraproject.org/) (Abe et al., 2020). The reference sequences used to design the primers were obtained from the Candid #1 vaccine strain (accession number: AY746354.1 for the L segment and AY746353.1 for the S segment). Viral RNA was amplified using PrimeScript II High Fidelity One Step RT-PCR Kit (Takara Bio, Shiga, Japan) under the following reaction conditions: 45 °C for 10 min, 94 °C for 2 min, 98 °C for 10 s, 55 °C for 15 s, and 68 °C for 10 s, for a total of 30 cycles. The products were then gel-purified using the QIAquick Gel Extraction Kit (Qiagen), according to the manufacturer’s instructions. Purified PCR products were sequenced using the BigDye Terminator v3.1 cycle sequencing kit (Thermo Fisher Scientific, MA, USA) and a ABI3500 sequencer (Thermo Fisher Scientific). Consensus sequences were generated and analyzed using GENETYX (GENETYX Corp., Tokyo, Japan) and SnapGene® softwares (GSL Biotech; available at snapgene.com). The sequences were submitted to the DNA Data Bank of Japan (DDBJ) (accession numbers: LC637306 for Candid1-P0-RdRp, LC637307 for Candid1-P0-Z, LC637308 for Candid1-P0-GPC, LC637309 for Candid1-P0-NP, LC637310 for Candid1-P11-RdRp, LC637311 for Candid1-P11-Z, LC637312 for Candid1-P11-GPC, and LC637313 for Candid1-P11-NP genes).

### Virus growth analysis

To compare growth kinetics of Candid #1 and Candid #1-res viruses, 293T cells were infected with each virus at an MOI of 0.1 in 24-well plates in duplicates. Infected cells were incubated on ice for 30 min with shaking the plate every 10 min. After inoculum removal, cells were washed twice with DMEM, and fresh media was added. Cells were incubated at 37 °C and viral titers were measured at 8, 12, 24, and 28 h post infection.

### Determination of the mutation frequency

Candid #1 and Candid #1-res viruses were used to infect 293T cells (MOI: 0.01) in the presence of 20 µM favipiravir or DMSO. At 48 hpi, RNA was extracted from culture supernatants and used to amplify a part of the NP gene using primer number five described in **Table S1**, with the high-fidelity One Step RT-PCR Kit (Takara Bio). PCR products were gel purified and cloned into the pCR4-TOPO vector using the Zero Blunt Topo Cloning Kit (Invitrogen). The clones were sequenced as described above. A fragment of 450 bp was used for nucleotide polymorphism analysis.

### Nucleoside supplementation assay

293T cells were infected with JUNV (MOI: 0.01). After adsorption at 37 °C for 1 h and removal of the inoculum, serial dilutions of the nucleosides adenosine (Sigma), guanosine (Sigma), thymine (Sigma), cytosine (Sigma), and uracil (Sigma) were added to the cells in combination with 50 µM (approximately 10 times the IC_50_) of favipiravir in triplicate. Cells treated with DMSO or favipiravir alone were used as controls. At 48 hpi, a plaque assay was performed to measure viral titers. Cells were visually inspected for any signs of cytotoxicity upon nucleoside treatment, and no toxic effects were observed. Results were expressed as a percentage reduction of the favipiravir anti-JUNV activity.

### Pseudotyped VSV production and virus entry assay

Full-length coding regions of JUNV GPC-A168 (from P0) and GPC-T168 (from P11) were cloned into the pCAGGS mammalian expression vector. Plasmids were designated as pC-GPC-A168 and pC-GPC-T168. Pseudotyped vesicular stomatitis virus (VSV) with a luciferase reporter gene, bearing JUNV GPC, was generated and titrated, as previously described (Kurosaki et al., 2018; Ushijima et al., 2021). Briefly, 293T cells were seeded in 6-well plates. After 8 h, cells were transfected with 3 µg of either each GPC expression plasmid or pCAGGS empty vector using TransIT LT-1 reagent (Mirus, Madison, WI, USA), according to the manufacturer’s instructions. At 24 hpt, cells were infected with G-complemented VSVΔG/Luc and incubated for 1 h at 37 °C for adsorption. The cells were washed three times with PBS, and DMEM containing 10% FBS was added to them. Pseudotyped viruses were collected at 24 hpi and labelled as Candid#1pv-A168 and Candid#1pv-T168, respectively. Viruses were stored at −80 °C until use. For the internalization assay, a confluent monolayer of 293T cells in a bottom-clear 96-well plate was cooled at 4 °C for 10 min and subsequently infected with either Candid#1pv-A168 or Candid#1pv-T168. The plates were further incubated at 4 °C for 30 min to allow the binding of viral particles to the receptor without initiation of the entry step (Carette et al., 2011). The cells were then washed three times with PBS to remove unbound viral particles. The plates were subsequently incubated at 37 ?. Luciferase activity was measured using the Steady-Glo Luciferase Assay System (Promega) and a TriStar LB 941 microplate reader (Berthord Japan K.K., Tokyo, Japan). Since there was a plateau effect at 24 hpi (data not shown), we considered the signal activity at this time point to be 100%.

### Western blotting

Supernatants containing pseudotyped virus were briefly cleared from debris by centrifugation. Ultracentrifugation was performed over a 20% sucrose cushion to pellet virion (60,000 rpm for 30 min at 4 ?). For the detection of intracellular proteins, cells were lysed using lysis buffer (1% NP-40, 50 mM Tris-HCl [pH 8.0], 62.5 mM EDTA, and 0.4% sodium deoxycholate). Prepared samples were analyzed by separation on either 12% (for VSV samples and actin) or 7.5% (for RdRp) sodium dodecyl sulphate–polyacrylamide gels through electrophoresis (SDS-PAGE) and western blotting (WB), as previously described (Zadeh et al., 2020). FLAG-tagged proteins, VSV M protein, or β-Actin were detected using mouse monoclonal primary antibodies against FLAG (M2, F1804, Sigma), VSV M (Kerafast, MA, USA), or β-actin (Sigma), respectively, and HRP-conjugated anti-mouse IgG secondary antibody (Sigma). The labelled proteins were then visualized using ECL prime (GE Healthcare) and LAS3000 (GE Healthcare), according to the manufacturer’s instructions. The results were quantified using Multi Gauge software (Fujifilm, Tokyo, Japan).

### Quantitative real time-polymerase chain reaction (qPCR)

Relative quantification of pseudotyped VSV was performed with a qPCR assay using forward (5’-TGTACATCGGAATGGCAGGG-3’) and reverse (5’-TGCCTTCACAGTGAGCATGATAC-3’) primers specific to the VSV M gene. One-Step TB Green PrimeScript PLUS RT-PCR Kit (Takara Bio) was used under the following conditions: 42 °C for 5 min, 95 °C for 5 s, and 60 °C for 34 s, for a total of 40 cycles using an ABI 7500 thermocycler (Applied Biosystems, Foster City, CA, USA). To quantify the encapsidated viral RNA copy numbers, the free RNA not associated with virions was removed from the samples using the Benzonase nuclease (Sigma), according to the manufacturer’s instructions, prior to viral RNA extraction. Standard RNA was synthesized from a partial region of the GPC gene using the forward (5’-TAATACGACTCACTATAGGGCCAACCTTTTTGCAGGAGGC-3’) and reverse (5’-AGCTTCTTCTGTGCAGGATCTTCCTGCAAGCGCTAGGAAT-3’) primers and the <u>T7 RNA polymerase</u> (Promega), as previously described (Pemba et al., 2019). The prepared RNA was then serially diluted using DEPC-treated water to obtain a standard curve ranging from 10^2^–10^13^ copies/mL. To quantify the nano-luciferase mRNA extracted from the minigenome assay, relative qPCR was performed using *GAPDH* expression as a control, as previously described (Zadeh et al., 2020), and specific primers targeting the nano-luciferase transcript (Forward, 5’-GGGAGGTGTGTCCAGTTTGT-3’ and reverse, 5’-CCGCTCAGACCTTCATACGG-3’).

### Minigenome assay

To compare the polymerase activities of RdRp-N462 and RdRp-D462, an MG system was constructed based on the Candid #1 S segment. First the coding region of Candid #1 RdRp was amplified using RNA extracted from P0 or P11 viruses and the PrimeScript II High Fidelity One Step RT-PCR Kit (Takara Bio). Kozak sequence, a FLAG tag (N-terminal), and a linker sequence (5’-GGTAGCGGCAGCGGTAGC-3’) were added through three additional PCR reactions using PrimeStar GXL DNA polymerase (Takara Bio). PCR products were gel-purified in each step, as described elsewhere. The entire fragment was then infused into a pCAGGS expression vector using an Infusion HD cloning kit (Takara Bio), according to the manufacturer’s instructions. The plasmid expressing JUNV NP, pC-Candid-NP, was kindly provided by Dr Juan C. de la Torre (Scripps Research Institute) (Emonet et al., 2011). To construct the MG plasmid, sequences of the untranslated regions (UTRs) of the Candid #1-S segment (based on the accession number AY746353) containing the 3′ UTR, 5′ UTR, and intergenic region in an antisense orientation were synthesized (GENEWIZ, NJ, USA). An additional G residue was added upstream of the 3′ UTR to enhance the efficiency of the system (Emonet et al., 2011). The synthesized fragment was then cloned into a pHH21 plasmid under the control of the human polymerase-I promoter (Neumann et al., 1999). The nano-luciferase (nluc) reporter gene was then inserted into the NP locus. Further details of the constructs can be provided upon request. To perform the assay, plasmids of Candid #1 NP, MG (with nluc reporter), and RdRp-N462 or RdRp-D462 were transfected into 293T cells at a 1:1:1 ratio using TransIT LT-1 (Mirus, Madison, WI). To normalize transfection efficiency, the pGL4.75 Renilla luciferase (Rluc) plasmid (Promega) was co-transfected. After 24 or 48 h, the cells were lysed and divided into two clear-bottom 96-well plates. Equal volumes of nano-Glo or Renilla-Glo (Promega) were added to measure nluc and Rluc independently. Polymerase activity was determined by the ratio of nluc/Rluc and expressed as relative luciferase induction.

### Drug combination assay and synergy analysis

To evaluate the combinational efficacy of favipiravir with either ribavirin or remdesivir, 293T cells infected with JUNV (MOI: 0.1) were treated with two-fold serially diluted combinations of the drugs at the indicated concentrations. Viral titers were determined at 48 hpi by plaque assay and represented as the percentage inhibition compared to DMSO control for each drug combination. Synergistic inhibition against JUNV growth was determined using SynergyFinder (https://synergyfinder.fimm.fi/) with the Zero Interaction Potency (ZIP) model as previously described (Imamura et al., 2021). A synergy score (δ-score) of less than −10 is considered as antagonistic, the score range of −10 to 10 suggests an additive drug interaction, and a score greater than 10 indicates a synergistic effect (Ianevski et al., 2020).

### Statistical analysis

Non-linear regression analysis was performed to analyze the dose-response of antivirals. The Mann–Whitney *U* rank test was used to compare the mutational frequency of viruses. Other statistical tests are mentioned in the respective figure legends. All analyses were performed using Prism version 8 (GraphPad Software Inc., La Jolla, CA, USA). Graphical representations were created using the web-based software, BioRender (https://biorender.com/).

## ABBREVIATIONS

JUNV: Junin virus
AHF: Argentine hemorrhagic fever
RdRp: RNA-dependent RNA polymerase
GPC: Glycoprotein precursor
VSV: vesicular stomatitis virus.

## AUTHOR APPROVALS

All authors have seen and approved the manuscript, and that it hasn’t been accepted or published elsewhere.

## ACKNOWLEDGMENTS

We are grateful to Dr. Haruka Abe for his helpful discussions and valuable comments on this manuscript. We are also grateful to Drs. Yasuteru Sakurai and Rokusuke Yoshikawa for their support and useful comments during this research work. We thank Dr. Juan C. de la Torre (Scripps Research Institute, California, USA) for providing the JUNV Candid #1 strain. We also thank Editage (www.editage.com) for English language editing. Favipiravir was kindly provided by FUJIFILM Toyama Chemical CO., LTD.

---

**Table S1.**
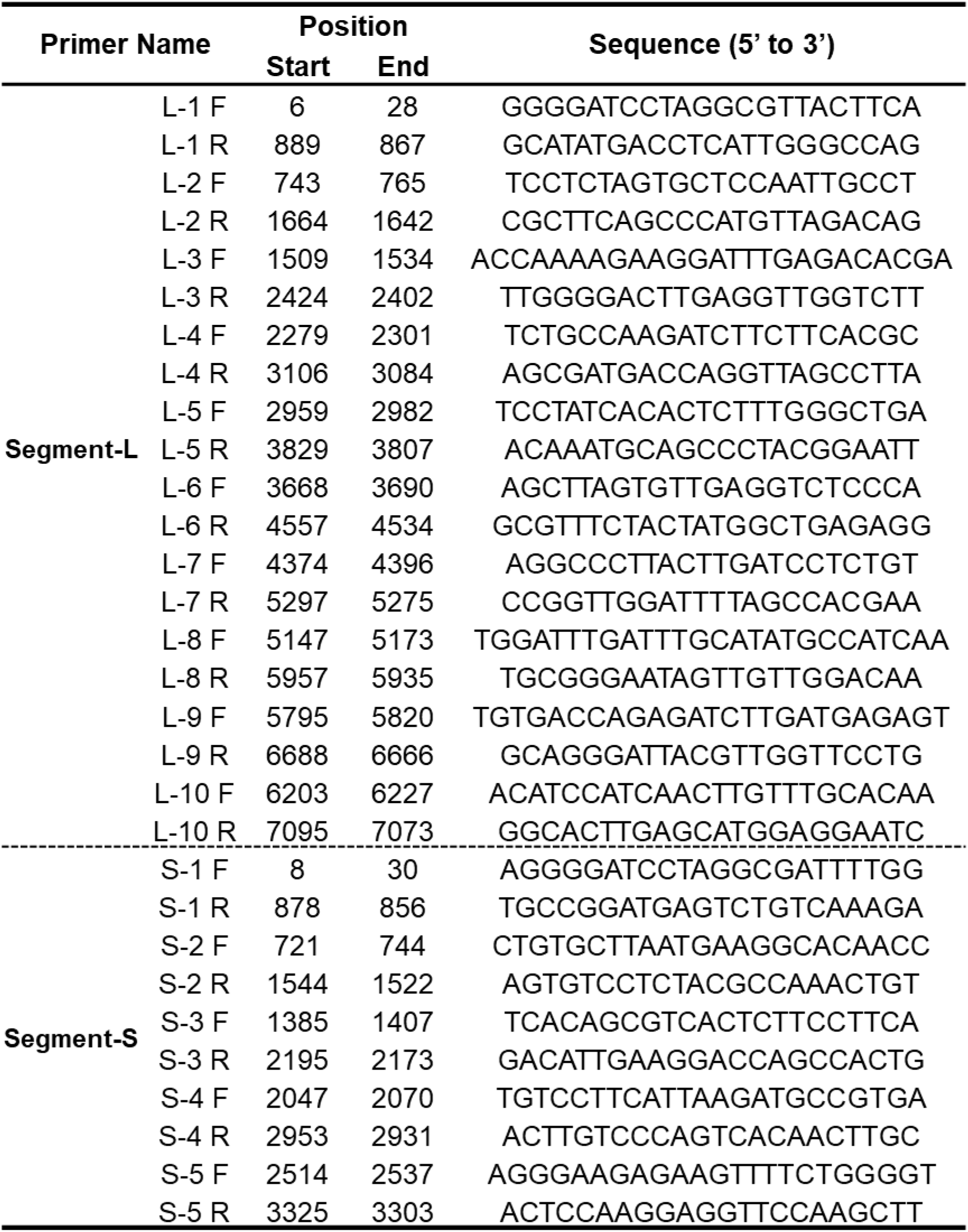
Primer list used for sequencing of JUNV genome L and S segments.

**Figure S1.**
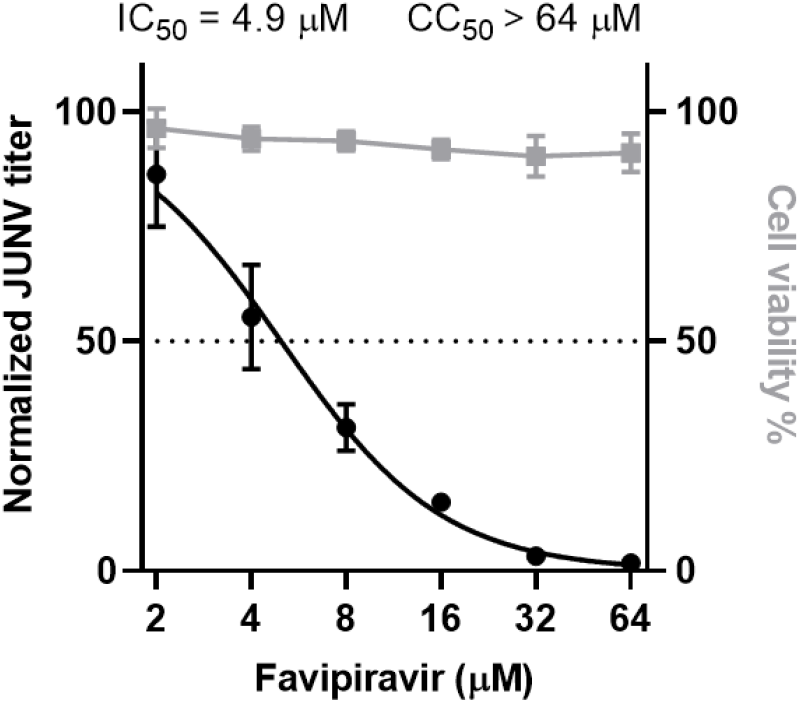
Determination of favipiravir IC50 value. 293T cells were infected with Candid #1 (MOI: 0.1). After adsorption, media containing serial dilutions of favipiravir was added. At 48 hpi, supernatant was collected and viral titers were determined by plaque assay. Error bars indicate ±SD; three independent experiments in duplicates (n = 6) were performed; nonlinear regression analysis was applied.

**Figure S2.**
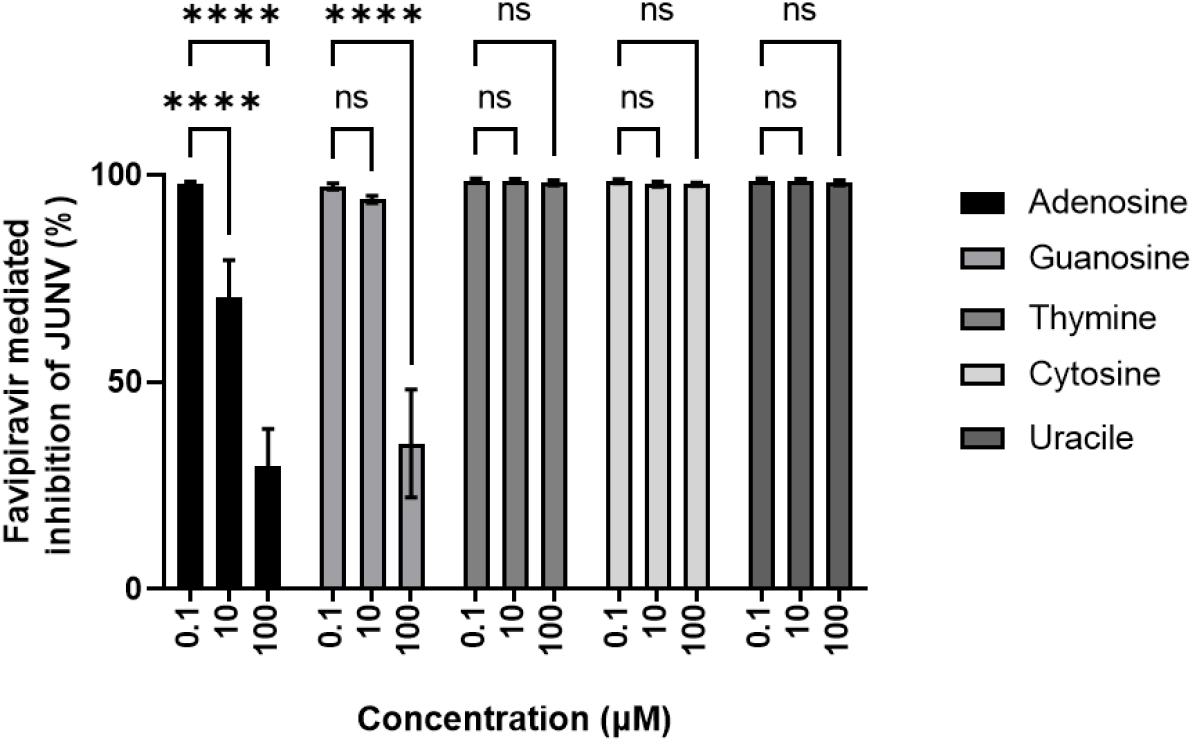
Nucleoside supplementation assay. 293T cells infected with JUNV (MOI: 0.01). were treated with serial dilutions of nucleosides adenosine, guanosine, thymine, cytosine, and uracil in combination with 50 µM of favipiravir. At 48 hpi, viral titers were measured by plaque assay. Titers were normalized to anti-JUNV activity of favipiravir to estimate the reversal imposed by nucleotide supplementations. Error bars indicate ±SD; two independent experiments in duplicates (n = 6) were performed. Statistical significance was determined by 2-way ANOVA tests (ns indicates not significant and *** indicates *P* < 0.001).

**Figure S3.**
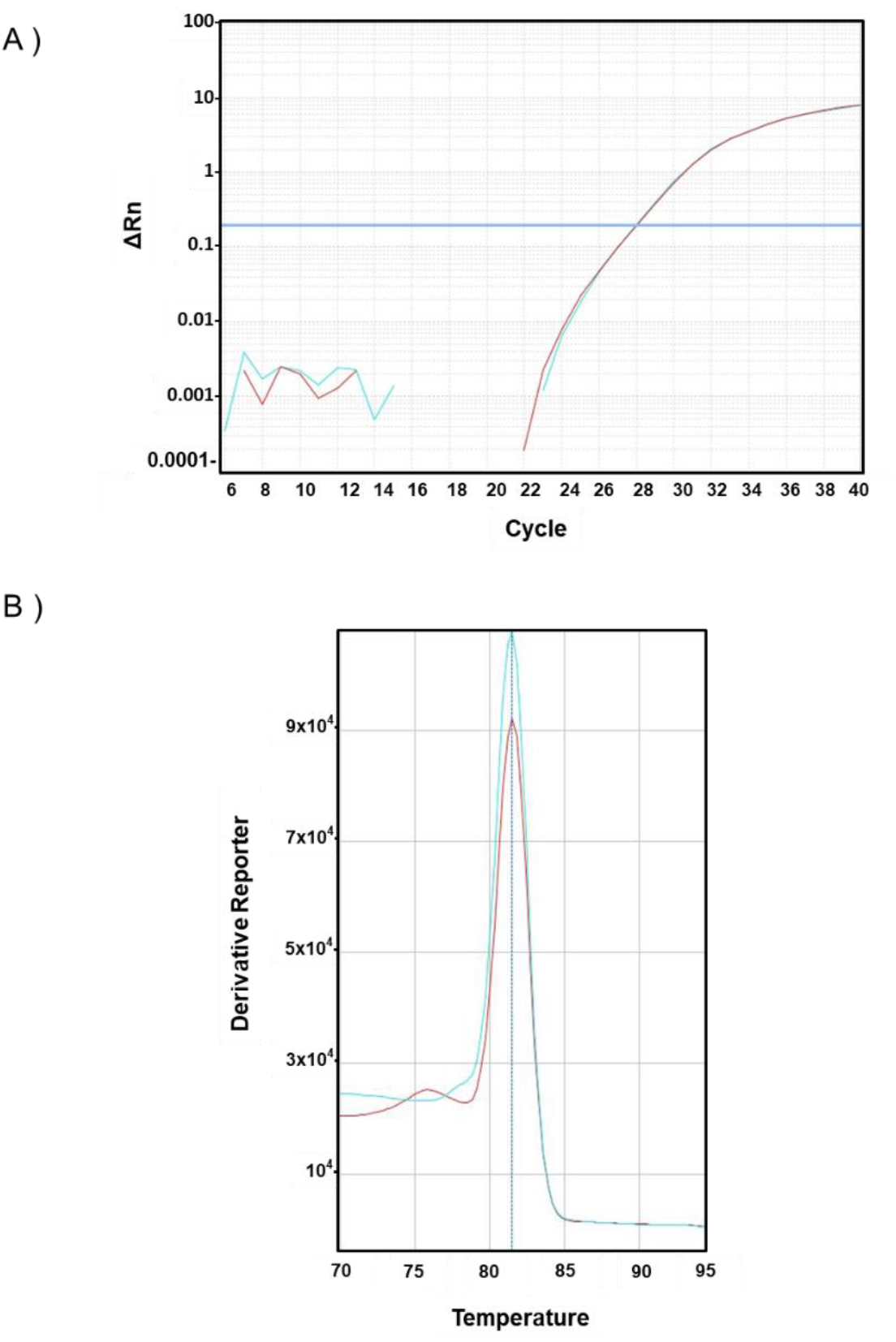
The amplification results of qPCR assay. (A) The amplification plot for VSV-M detection from Candid #1pv-A168 (red) or Candid #1pv-T168 (blue) showing a CT value of 27.99 and 27.95 are shown respectively. (B) Melting curve analysis at the end of the amplification run confirms the specificity of the assay at the end of the amplification.

**Figure S4.**
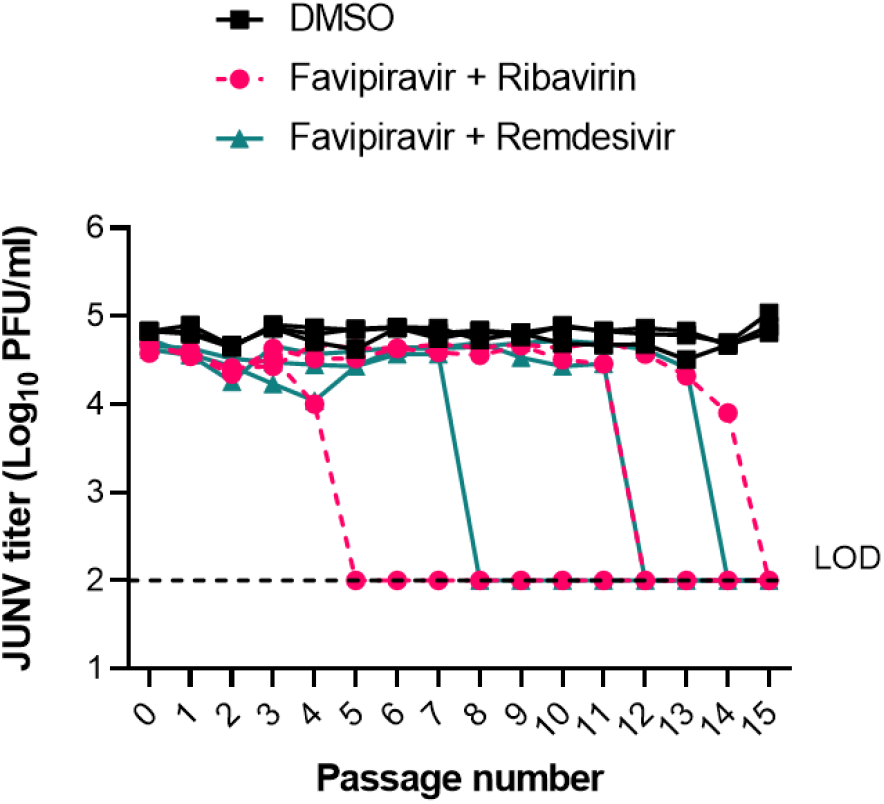
Serial passaging of JUNV treated with combination of favipiravir and ribavirin or remdesivir. 293T cells were infected with JUNV at MOI of 0.01 for initial inoculation and 10-fold dilutions for the remaining passages (n = 3). After adsorption, cells were treated with combinations of favipiravir (0.3 µM), ribavirin (0.3 µM), and remdesivir (1 nM), and titers were determined at 48 hpi by plaque assay.

**Figure S5.**
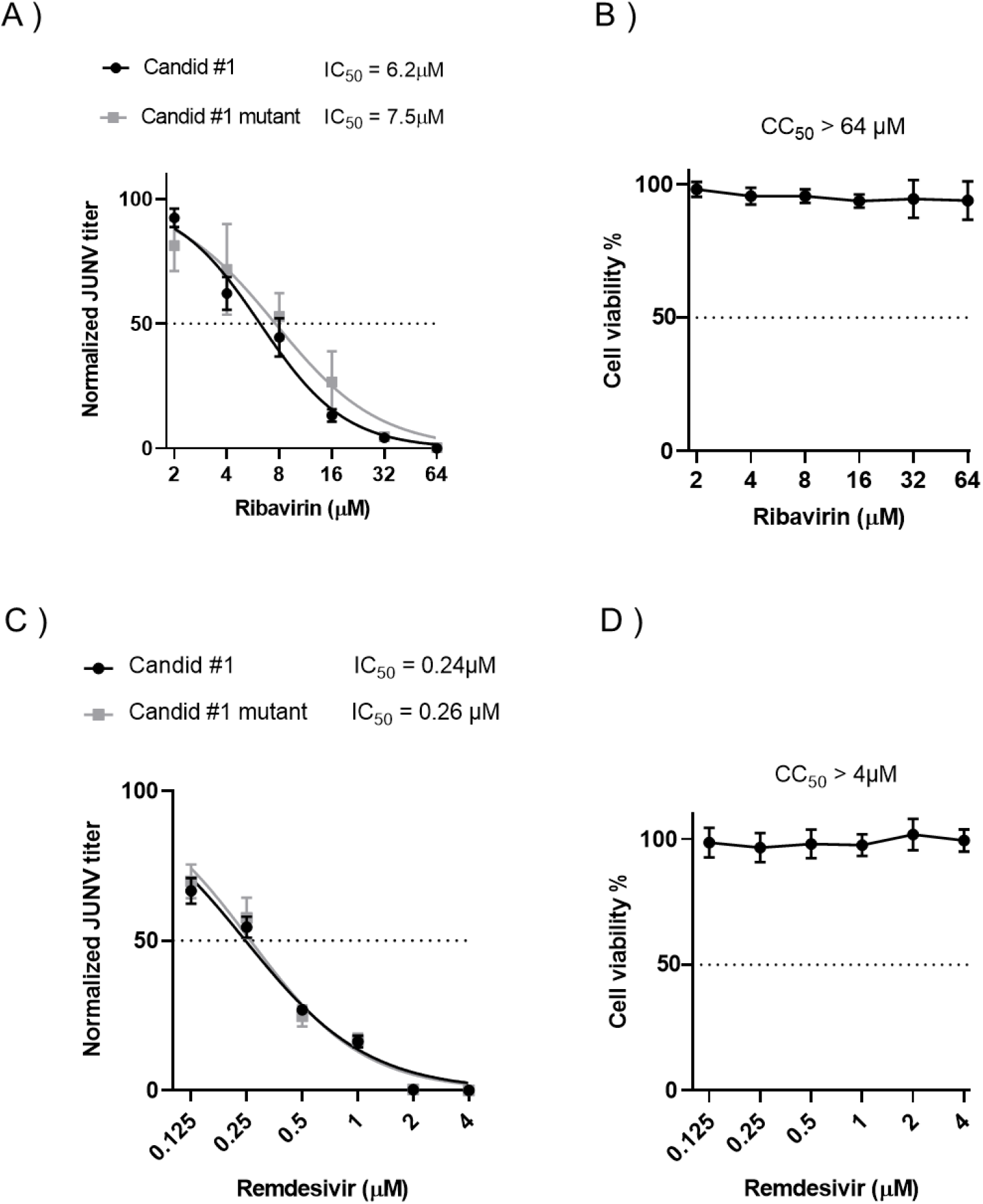
JUNV Candid #1-mutant virus remains susceptible to ribavirin and remdesivir. 293T cell were infected with JUNV Candid #1 or Candid #1-mutant virus (MOI: 0.1). Media containing the indicated concentrations of ribavirin was added. At 48 hpi, viral titers were measured by plaques assay. Cytotoxicity assay was performed as described in materials and methods. Error bars indicate ±SD; three independent experiments in duplicate (n = 6) were performed; nonlinear regression analysis was applied.

## REFERENCES

Abe, H., Ushijima, Y., Bikangui, R., Ondo, G.N., Zadeh, V.R., Pemba, C.M., Mpingabo, P.I., Igasaki, Y., Vries S.G. de, Grobusch, M.P., Loembe, M.M., Agnandji, S.T., Lell, B., Yasuda, J., 2020. First evidence for continuous circulation of hepatitis A virus subgenotype IIA in Central Africa. J. Viral Hepat. 27, 1234–1242. https://doi.org/10.1111/jvh.13348

Arias, A., Thorne, L., Goodfellow, I., 2014. Favipiravir elicits antiviral mutagenesis during virus replication in vivo. eLife 3. https://doi.org/10.7554/eLife.03679

Ávila, A.I. de, Gallego, I., Soria, M.E., Gregori, J., Quer, J., Esteban, J.I., Rice, C.M., Domingo, E., Perales, C., 2016. Lethal Mutagenesis of Hepatitis C Virus Induced by Favipiravir. PLOS ONE 11, e0164691. https://doi.org/10.1371/journal.pone.0164691

Bassi, M.R., Sempere, R.N., Meyn, P., Polacek, C., Arias, A., 2018. Extinction of Zika Virus and Usutu Virus by Lethal Mutagenesis Reveals Different Patterns of Sensitivity to Three Mutagenic Drugs. Antimicrob. Agents Chemother. 62. https://doi.org/10.1128/AAC.00380-18

Borio, L., Inglesby, T., Peters, C.J., Schmaljohn, A.L., Hughes, J.M., Jahrling, P.B., Ksiazek, T., Johnson, K.M., Meyerhoff, A., O’Toole, T., Ascher, M.S., Bartlett, J., Breman, J.G., Eitzen, E.M., Hamburg, M., Hauer, J., Henderson, D.A., Johnson, R.T., Kwik, G., Layton, M., Lillibridge, S., Nabel, G.J., Osterholm, M.T., Perl, T.M., Russell, P., Tonat, K., Working Group on Civilian Biodefense, 2002. Hemorrhagic fever viruses as biological weapons: medical and public health management. JAMA 287, 2391–2405. https://doi.org/10.1001/jama.287.18.2391

Bruenn, J.A., 2003. A structural and primary sequence comparison of the viral RNA-dependent RNA polymerases. Nucleic Acids Res. 31, 1821–1829.

Brunotte, L., Lelke, M., Hass, M., Kleinsteuber, K., Becker-Ziaja, B., Günther, S., 2011. Domain Structure of Lassa Virus L Protein. J. Virol. 85, 324–333. https://doi.org/10.1128/JVI.00721-10

Carette, J.E., Raaben, M., Wong, A.C., Herbert, A.S., Obernosterer, G., Mulherkar, N., Kuehne, A.I., Kranzusch, P.J., Griffin, A.M., Ruthel, G., Cin, P.D., Dye, J.M., Whelan, S.P., Chandran, K., Brummelkamp, T.R., 2011. Ebola virus entry requires the cholesterol transporter Niemann– Pick C1. Nature 477, 340–343. https://doi.org/10.1038/nature10348

Carrillo-Bustamante, P., Nguyen, T.H.T., Oestereich, L., Günther, S., Guedj, J., Graw, F., 2017. Determining Ribavirin’s mechanism of action against Lassa virus infection. Sci. Rep. 7, 11693. https://doi.org/10.1038/s41598-017-10198-0

Cheung, P.P.H., Watson, S.J., Choy, K.-T., Sia, S.F., Wong, D.D.Y., Poon, L.L.M., Kellam, P., Guan, Y., Peiris, J.S.M., Yen, H.-L., 2014. Generation and characterization of influenza A viruses with altered polymerase fidelity. Nat. Commun. 5, 1–13. https://doi.org/10.1038/ncomms5794

Delang, L., Abdelnabi, R., Neyts, J., 2018. Favipiravir as a potential countermeasure against neglected and emerging RNA viruses. Antiviral Res. 153, 85–94. https://doi.org/10.1016/j.antiviral.2018.03.003

Delang, L., Segura Guerrero, N., Tas, A., Quérat, G., Pastorino, B., Froeyen, M., Dallmeier, K., Jochmans, D., Herdewijn, P., Bello, F., Snijder, E.J., de Lamballerie, X., Martina, B., Neyts, J., van Hemert, M.J., Leyssen, P., 2014. Mutations in the chikungunya virus non-structural proteins cause resistance to favipiravir (T-705), a broad-spectrum antiviral. J. Antimicrob. Chemother. 69, 2770–2784. https://doi.org/10.1093/jac/dku209

Emonet, S.E., Urata, S., de la Torre, J.C., 2011. Arenavirus reverse genetics: New approaches for the investigation of arenavirus biology and development of antiviral strategies. Virology, Special Reviews Issue 2011 411, 416–425. https://doi.org/10.1016/j.virol.2011.01.013

Emonet, S.F., Seregin, A.V., Yun, N.E., Poussard, A.L., Walker, A.G., de la Torre, J.C., Paessler, S., 2011. Rescue from Cloned cDNAs and In Vivo Characterization of Recombinant Pathogenic Romero and Live-Attenuated Candid #1 Strains of Junin Virus, the Causative Agent of Argentine Hemorrhagic Fever Disease. J. Virol. 85, 1473–1483. https://doi.org/10.1128/JVI.02102-10

Enria, D.A., Briggiler, A.M., Sánchez, Z., 2008. Treatment of Argentine hemorrhagic fever. Antiviral Res., Special Issue: Treatment of highly pathogenic RNA viral infections 78, 132–139. https://doi.org/10.1016/j.antiviral.2007.10.010

Espy, N., Nagle, E., Pfeffer, B., Garcia, K., Chitty, A.J., Wiley, M., Sanchez-Lockhart, M., Bavari, S., Warren, T., Palacios, G., 2019. T-705 induces lethal mutagenesis in Ebola and Marburg populations in macaques. Antiviral Res. 170, 104529. https://doi.org/10.1016/j.antiviral.2019.06.001

Feld, J.J., Hoofnagle, J.H., 2005. Mechanism of action of interferon and ribavirin in treatment of hepatitis C. Nature 436, 967–972. https://doi.org/10.1038/nature04082

Furuta, Y., Takahashi, K., Kuno-Maekawa, M., Sangawa, H., Uehara, S., Kozaki, K., Nomura, N., Egawa, H., Shiraki, K., 2005. Mechanism of action of T-705 against influenza virus. Antimicrob. Agents Chemother. 49, 981–986. https://doi.org/10.1128/AAC.49.3.981-986.2005

Goldhill, D.H., Langat, P., Xie, H., Galiano, M., Miah, S., Kellam, P., Zambon, M., Lackenby, A., Barclay, W.S., 2018a. Determining the Mutation Bias of Favipiravir in Influenza Virus Using Next-Generation Sequencing. J. Virol. 93, e01217–18, /jvi/93/2/JVI.01217-18.atom. https://doi.org/10.1128/JVI.01217-18

Goldhill, D.H., Velthuis, A.J.W. te, Fletcher, R.A., Langat, P., Zambon, M., Lackenby, A., Barclay, W.S., 2018b. The mechanism of resistance to favipiravir in influenza. Proc. Natl. Acad. Sci. 115, 11613–11618. https://doi.org/10.1073/pnas.1811345115

Gowen, B.B., Hickerson, B.T., York, J., Westover, J.B., Sefing, E.J., Bailey, K.W., Wandersee, L., Nunberg, J.H., 2021. Second-generation live-attenuated Candid#1 vaccine virus resists reversion and protects against lethal Junín virus infection in guinea pigs. J. Virol. https://doi.org/10.1128/JVI.00397-21

Gowen, B.B., Juelich, T.L., Sefing, E.J., Brasel, T., Smith, J.K., Zhang, L., Tigabu, B., Hill, T.E., Yun, T., Pietzsch, C., Furuta, Y., Freiberg, A.N., 2013. Favipiravir (T-705) Inhibits Junín Virus Infection and Reduces Mortality in a Guinea Pig Model of Argentine Hemorrhagic Fever. PLoS Negl. Trop. Dis. 7, e2614. https://doi.org/10.1371/journal.pntd.0002614

Gowen, B.B., Westover, J.B., Sefing, E.J., Van Wettere, A.J., Bailey, K.W., Wandersee, L., Komeno, T., Furuta, Y., 2017. Enhanced protection against experimental Junin virus infection through the use of a modified favipiravir loading dose strategy. Antiviral Res. 145, 131–135. https://doi.org/10.1016/j.antiviral.2017.07.019

Gowen, B.B., Wong, M.-H., Jung, K.-H., Sanders, A.B., Mendenhall, M., Bailey, K.W., Furuta, Y., Sidwell, R.W., 2007. In Vitro and In Vivo Activities of T-705 against Arenavirus and Bunyavirus Infections. Antimicrob. Agents Chemother. 51, 3168–3176. https://doi.org/10.1128/AAC.00356-07

Grande-Pérez, A., Martin, V., Moreno, H., de la Torre, J.C., 2015. Arenavirus Quasispecies and Their Biological Implications. Quasispecies Theory Exp. Syst. 392, 231–275. https://doi.org/10.1007/82_2015_468

Guedj, J., Piorkowski, G., Jacquot, F., Madelain, V., Nguyen, T.H.T., Rodallec, A., Gunther, S., Carbonnelle, C., Mentré, F., Raoul, H., Lamballerie, X. de, 2018. Antiviral efficacy of favipiravir against Ebola virus: A translational study in cynomolgus macaques. PLOS Med. 15, e1002535. https://doi.org/10.1371/journal.pmed.1002535

Ianevski, A., Giri, A.K., Aittokallio, T., 2020. SynergyFinder 2.0: visual analytics of multi-drug combination synergies. Nucleic Acids Res. 48, W488–W493. https://doi.org/10.1093/nar/gkaa216

Imamura, K., Sakurai, Y., Enami, T., Shibukawa, R., Nishi, Y., Ohta, A., Shu, T., Kawaguchi, J., Okada, S., Hoenen, T., Yasuda, J., Inoue, H., 2021. iPSC screening for drug repurposing identifies anti-RNA virus agents modulating host cell susceptibility. FEBS Open Bio 11, 1452–1464. https://doi.org/10.1002/2211-5463.13153

Kenyon, R.H., Canonico, P.G., Green, D.E., Peters, C.J., 1986. Effect of ribavirin and tributylribavirin on argentine hemorrhagic fever (Junin virus) in guinea pigs. Antimicrob. Agents Chemother. 29, 521–523. https://doi.org/10.1128/AAC.29.3.521

Kurosaki, Y., Ueda, M.T., Nakano, Y., Yasuda, J., Koyanagi, Y., Sato, K., Nakagawa, S., 2018. Different effects of two mutations on the infectivity of Ebola virus glycoprotein in nine mammalian species. J. Gen. Virol. 99, 181–186. https://doi.org/10.1099/jgv.0.000999

Lingas, G., Rosenke, K., Safronetz, D., Guedj, J., 2021. Lassa viral dynamics in non-human primates treated with favipiravir or ribavirin. PLOS Comput. Biol. 17, e1008535. https://doi.org/10.1371/journal.pcbi.1008535

Lo, M.K., Albariño, C.G., Perry, J.K., Chang, S., Tchesnokov, E.P., Guerrero, L., Chakrabarti, A., Shrivastava-Ranjan, P., Chatterjee, P., McMullan, L.K., Martin, R., Jordan, R., Götte, M., Montgomery, J.M., Nichol, S.T., Flint, M., Porter, D., Spiropoulou, C.F., 2020. Remdesivir targets a structurally analogous region of the Ebola virus and SARS-CoV-2 polymerases. Proc. Natl. Acad. Sci. 117, 26946–26954. https://doi.org/10.1073/pnas.2012294117

Madelain, V., Guedj, J., Mentré, F., Nguyen, T.H.T., Jacquot, F., Oestereich, L., Kadota, T., Yamada, K., Taburet, A.-M., Lamballerie, X. de, Raoul, H., 2017. Favipiravir Pharmacokinetics in Nonhuman Primates and Insights for Future Efficacy Studies of Hemorrhagic Fever Viruses. Antimicrob. Agents Chemother. 61. https://doi.org/10.1128/AAC.01305-16

McKee, K.T., Huggins, J.W., Trahan, C.J., Mahlandt, B.G., 1988. Ribavirin prophylaxis and therapy for experimental argentine hemorrhagic fever. Antimicrob. Agents Chemother. 32, 1304–1309. https://doi.org/10.1128/AAC.32.9.1304

McKee, K.T., Oro, J.G., Kuehne, A.I., Spisso, J.A., Mahlandt, B.G., 1993. Safety and immunogenicity of a live-attenuated Junin (Argentine hemorrhagic fever) vaccine in rhesus macaques. Am. J. Trop. Med. Hyg. 48, 403–411. https://doi.org/10.4269/ajtmh.1993.48.403

Mendenhall, M., Russell, A., Juelich, T., Messina, E.L., Smee, D.F., Freiberg, A.N., Holbrook, M.R., Furuta, Y., de la Torre, J.-C., Nunberg, J.H., Gowen, B.B., 2011a. T-705 (Favipiravir) Inhibition of Arenavirus Replication in Cell Culture. Antimicrob. Agents Chemother. 55, 782–787. https://doi.org/10.1128/AAC.01219-10

Mendenhall, M., Russell, A., Smee, D.F., Hall, J.O., Skirpstunas, R., Furuta, Y., Gowen, B.B., 2011b. Effective Oral Favipiravir (T-705) Therapy Initiated after the Onset of Clinical Disease in a Model of Arenavirus Hemorrhagic Fever. PLoS Negl. Trop. Dis. 5, e1342. https://doi.org/10.1371/journal.pntd.0001342

Neagu, I.A., Olejarz, J., Freeman, M., Rosenbloom, D.I.S., Nowak, M.A., Hill, A.L., 2018. Life cycle synchronization is a viral drug resistance mechanism. PLOS Comput. Biol. 14, e1005947. https://doi.org/10.1371/journal.pcbi.1005947

Neumann, G., Watanabe, T., Ito, H., Watanabe, S., Goto, H., Gao, P., Hughes, M., Perez, D.R., Donis, R., Hoffmann, E., Hobom, G., Kawaoka, Y., 1999. Generation of influenza A viruses entirely from cloned cDNAs. Proc. Natl. Acad. Sci. 96, 9345–9350. https://doi.org/10.1073/pnas.96.16.9345

NIAID Emerging Infectious Diseases/Pathogens | NIH: National Institute of Allergy and Infectious Diseases [WWW Document], n.d. URL https://www.niaid.nih.gov/research/emerging-infectious-diseases-pathogens (accessed 11.24.18).

Pauly, M.D., Lauring, A.S., 2015. Effective Lethal Mutagenesis of Influenza Virus by Three Nucleoside Analogs. J. Virol. 89, 3584–3597. https://doi.org/10.1128/JVI.03483-14

Pemba, C.M., Kurosaki, Y., Yoshikawa, R., Oloniniyi, O.K., Urata, S., Sueyoshi, M., Zadeh, V.R., Nwafor, I., Iroezindu, M.O., Ajayi, N.A., Chukwubike, C.M., Chika-Igwenyi, N.M., Ndu, A.C., Nwidi, D.U., Maehira, Y., Unigwe, U.S., Ojide, C.K., Onwe, E.O., Yasuda, J., 2019. Development of an RT-LAMP assay for the detection of Lassa viruses in southeast and south-central Nigeria. J. Virol. Methods 269, 30–37. https://doi.org/10.1016/j.jviromet.2019.04.010

Peng, R., Xu, X., Jing, J., Wang, M., Peng, Q., Liu, S., Wu, Y., Bao, X., Wang, P., Qi, J., Gao, G.F., Shi, Y., 2020. Structural insight into arenavirus replication machinery. Nature 579, 615–619. https://doi.org/10.1038/s41586-020-2114-2

Pfeiffer, J.K., Kirkegaard, K., 2005. Increased Fidelity Reduces Poliovirus Fitness and Virulence under Selective Pressure in Mice. PLOS Pathog. 1, e11. https://doi.org/10.1371/journal.ppat.0010011

Pfeiffer, J.K., Kirkegaard, K., 2003. A single mutation in poliovirus RNA-dependent RNA polymerase confers resistance to mutagenic nucleotide analogs via increased fidelity. Proc. Natl. Acad. Sci. 100, 7289–7294. https://doi.org/10.1073/pnas.1232294100

Rosenke, K., Feldmann, H., Westover, J.B., Hanley, P.W., Martellaro, C., Feldmann, F., Saturday, G., Lovaglio, J., Scott, D.P., Furuta, Y., Komeno, T., Gowen, B.B., Safronetz, D., 2018. Use of Favipiravir to Treat Lassa Virus Infection in Macaques. Emerg. Infect. Dis. 24, 1696–1699. https://doi.org/10.3201/eid2409.180233

Sadeghipour, S., Bek, E.J., McMinn, P.C., 2013. Ribavirin-Resistant Mutants of Human Enterovirus 71 Express a High Replication Fidelity Phenotype during Growth in Cell Culture. J. Virol. 87, 11.

Sedaghat, A.R., Wilke, C.O., 2011. Kinetics of the viral cycle influence pharmacodynamics of antiretroviral therapy. Biol. Direct 6, 42. https://doi.org/10.1186/1745-6150-6-42

Stephan, B.I., Lozano, M.E., Goñi, S.E., 2013. Watching Every Step of the Way: Junín Virus Attenuation Markers in the Vaccine Lineage. Curr. Genomics 14, 415–424. https://doi.org/10.2174/138920291407131220153526

Suemori, K., Saijo, M., Yamanaka, A., Himeji, D., Kawamura, M., Haku, T., Hidaka, M., Kamikokuryo, C., Kakihana, Y., Azuma, T., Takenaka, K., Takahashi, T., Furumoto, A., Ishimaru, T., Ishida, M., Kaneko, M., Kadowaki, N., Ikeda, K., Sakabe, S., Taniguchi, T., Ohge, H., Kurosu, T., Yoshikawa, T., Shimojima, M., Yasukawa, M., 2021. A multicenter non-randomized, uncontrolled single arm trial for evaluation of the efficacy and the safety of the treatment with favipiravir for patients with severe fever with thrombocytopenia syndrome. PLoS Negl. Trop. Dis. 15, e0009103. https://doi.org/10.1371/journal.pntd.0009103

Szemiel, A.M., Merits, A., Orton, R.J., MacLean, O., Pinto, R.M., Wickenhagen, A., Lieber, G., Turnbull, M.L., Wang, S., Mair, D., Filipe, A. da S., Willett, B.J., Wilson, S.J., Patel, A.H., Thomson, E.C., Palmarini, M., Kohl, A., Stewart, M.E., 2021. In vitro evolution of Remdesivir resistance reveals genome plasticity of SARS-CoV-2. bioRxiv 2021.02.01.429199. https://doi.org/10.1101/2021.02.01.429199

Tchesnokov, E.P., Gordon, C.J., Woolner, E., Kocincova, D., Perry, J.K., Feng, J.Y., Porter, D.P., Gotte, M., 2020. Template-dependent inhibition of coronavirus RNA-dependent RNA polymerase by remdesivir reveals a second mechanism of action. J. Biol. Chem. https://doi.org/10.1074/jbc.AC120.015720

Urata, S., Yasuda, J., 2012. Molecular Mechanism of Arenavirus Assembly and Budding. Viruses 4, 2049–2079. https://doi.org/10.3390/v4102049

Ushijima, Y., Abe, H., Ozeki, T., Ondo, G.N., Mbadinga, M.J.V.M., Bikangui, R., Nze-Nkogue, C., Akomo-Okoue, E.F., Ella, G.W.E., Koumba, L.B.M., Nso, B.C.B.B., Mintsa-Nguema, R., Makouloutou-Nzassi, P., Makanga, B.K., Nguelet, F.L.M., Zadeh, V.R., Urata, S., Mbouna, A.V.N., Loembe, M.M., Agnandji, S.T., Lell, B., Yasuda, J., 2021. Identification of potential novel hosts and the risk of infection of lymphocytic choriomeningitis virus in humans in Gabon, Central Africa. Int. J. Infect. Dis. https://doi.org/10.1016/j.ijid.2021.02.105

Veliziotis, I., Roman, A., Martiny, D., Schuldt, G., Claus, M., Dauby, N., Wijngaert, S.V. den, Martin, C., Nasreddine, R., Perandones, C., Mahieu, R., Swaan, C., Praet, S.V., Konopnicki, D., Morales, M.A., Malvy, D., Stevens, E., Dechamps, P., Vlieghe, E., Vandenberg, O., Günther, S., Gérard, M., n.d. Clinical Management of Argentine Hemorrhagic Fever using Ribavirin and Favipiravir, Belgium, 2020 - Volume 26, Number 7—July 2020 - Emerging Infectious Diseases journal - CDC. https://doi.org/10.3201/eid2607.200275

Vignuzzi, M., Stone, J.K., Arnold, J.J., Cameron, C.E., Andino, R., 2006. Quasispecies diversity determines pathogenesis through cooperative interactions in a viral population. Nature 439, 344–348. https://doi.org/10.1038/nature04388

Wang, Y., Li, G., Yuan, S., Gao, Q., Lan, K., Altmeyer, R., Zou, G., 2016. In Vitro Assessment of Combinations of Enterovirus Inhibitors against Enterovirus 71. Antimicrob. Agents Chemother. 60, 5357–5367. https://doi.org/10.1128/AAC.01073-16

Weissenbacher, M.C., Calello, M.A., Merani, M.S., Rodriguez, M., McCormick, J.B., 1986. Therapeutic Effect of the Antiviral Agent Ribavirin in Junin Virus Infection of Primates. J. Med. Virol. 20, 261–267. https://doi.org/10.1002/jmv.1890200308

Westover, J.B., Sefing, E.J., Bailey, K.W., Van Wettere, A.J., Jung, K.-H., Dagley, A., Wandersee, L., Downs, B., Smee, D.F., Furuta, Y., Bray, M., Gowen, B.B., 2016. Low-dose ribavirin potentiates the antiviral activity of favipiravir against hemorrhagic fever viruses. Antiviral Res. 126, 62–68. https://doi.org/10.1016/j.antiviral.2015.12.006

Zadeh, V.R., Urata, S., Sakaguchi, M., Yasuda, J., 2020. Human BST-2/tetherin inhibits Junin virus release from host cells and its inhibition is partially counteracted by viral nucleoprotein. J. Gen. Virol. https://doi.org/10.1099/jgv.0.001414

Ziegler, C.M., Botten, J.W., 2020. Defective Interfering Particles of Negative-Strand RNA Viruses. Trends Microbiol. 28, 554–565. https://doi.org/10.1016/j.tim.2020.02.006

